# A RecBCD-like enzyme regulates expression of bacterial defense operons in response to DNA damage

**DOI:** 10.64898/2026.10.31.763320

**Authors:** Lydia R. Chambers, Kishen M. Patel, Minheng Gong, Dwaipayan Basu, Elina Tian, Stephanie N. Ouderkirk, Yaya Zhou, Qiaozhen Ye, Brian Q. Pham, Tatiana V. Mishanina, Aaron T. Whiteley, Kevin D. Corbett

## Abstract

Bacteria must quickly respond to stress signals including bacteriophage (phage) infection, invasion by mobile genetic elements, and other threats. DNA damage is a stress-signaling nexus in bacteria, initiating global responses such as the SOS response and triggering expression of specific defense and immune pathways through DNA damage-sensing transcriptional regulators. Here we identify a three-protein pathway termed Cerberus that senses dsDNA ends that become available during DNA damage or inhibition of the key DNA repair complex RecBCD, and controls expression of an associated defense/immune operon. The central player in Cerberus is a large peptidase-helicase-nuclease fusion protein, CerC, that is architecturally similar to the RecBC complex itself. Upon DNA damage or RecBCD inhibition, CerC binds and unwinds free DNA ends to allosterically activate its fused HExxH metallopeptidase domain. CerC specifically cleaves the adjacently encoded anti-sigma factor CerB to release its cognate sigma factor CerA, allowing CerA to mediate transcription of a defense/immune operon encoded nearby. Cerberus therefore represents a pathway that has co-opted a key DNA repair machine for DNA damage sensing, and linked that activity to transcriptional control through anti-sigma factor cleavage.

## INTRODUCTION

Bacteria are under constant pressure from external threats including bacteriophages (phages), conjugating plasmids, and toxins produced by other bacteria. Through this struggle for survival, bacteria have evolved to encode a broad arsenal of defensive pathways with diverse mechanisms of action^1,2^. Operons encoding these pathways are typically clustered in genomic regions termed “defense islands,” which share characteristics with mobile genetic elements that enable these pathways to be transferred between bacteria via horizontal gene transfer^3–5^. Among these defensive pathways are anti-phage immune pathways, which commonly operate by directly targeting and destroying the DNA of an invading phage (termed direct defense), depleting key metabolites to slow phage replication, or triggering cell death to prevent phage replication (termed abortive infection)^6–10^. Because abortive infection pathways in particular are inherently costly or destructive, their activation must be tightly regulated.

Immune pathway regulation occurs through a range of mechanisms. Abortive infection pathways typically have an internal regulation mechanism, activating their toxic effectors only when infection is sensed through the presence of a particular molecular species (e.g., a phage capsid protein) or through metabolic changes (e.g., depletion of ATP or other nucleotides)^9,11^. In addition to these internal regulation mechanisms, many pathways are also regulated at the level of transcription. For example, some CRISPR-Cas systems utilize quorum sensing, allowing cells to coordinate defense activation based on cell density—which can be considered a proxy for phage-infection risk^12,13^. Many type II Restriction-Modification systems encode a controller (C) protein that fine-tunes the relative expression timing of the methylase and nuclease components to prevent destructive nuclease activity^14^. More directly, the expression of some pathways is controlled by dedicated transcription factors, including CapP-CapH^15^, BrxR/CapW^16–18^, and CapK-CapS^19^, that are encoded adjacent to or within the immune operon itself. Each of these regulators directly activates transcription of an associated immune operon in response to DNA damage by sensing the formation of single-stranded DNA (ssDNA), a byproduct of several DNA repair pathways^15–20^. The role of these transcriptional regulators is not fully understood, but they may enable bacteria to mount a fast and specific immune/defensive response, while avoiding the fitness cost of continual expression and possible aberrant activation of these potentially destructive pathways.

Here, we describe a three-gene operon, which we term Cerberus, that regulates transcription of an associated operon through a previously undescribed mechanism. The Cerberus operon includes a sigma factor–anti-sigma factor pair (CerA and CerB) and a multidomain peptidase-helicase-nuclease fusion protein (CerC). CerC is structurally related to well-characterized DNA repair-associated helicase-nuclease protein complexes including RecBCD, AddAB, and AdnAB, but where these proteins assemble into obligate heterodimers (or heterotrimers, as in RecBCD), CerC functions as a monomer. Moreover, CerC displays a distinctive domain organization including an N-terminal IrrE-like zinc metallopeptidase domain. We show that CerC binds and unwinds double-stranded DNA, generating a single-stranded tail that activates its own peptidase domain. CerC specifically cleaves the anti-sigma factor CerB, releasing the sigma factor CerA to drive expression of the associated operon. In *E. coli*, we find that Cerberus regulates expression of its associated defense/immune operon in response to either inhibition of RecBCD or the introduction of DNA damage that overwhelms RecBCD’s repair capacity. Thus, unlike other DNA damage-sensing transcriptional regulators, Cerberus integrates sensing of multiple stress signals to regulate defense/immune pathway expression.

## RESULTS

### Identification of *cerABC* operons associated with defensive pathways

Our recent work on bacterial ubiquitination-like (Bub) pathways revealed that a large fraction of Bub-encoding operons co-occur with DNA damage-sensing transcriptional regulators, including CapP-CapH (32% of Bub operons), CapW (8%), and CapK-CapS (11%)^21,22^. While manually inspecting the genomic neighborhoods of the remaining Bub operons, we found that an additional 21% co-occur with a three-gene operon, which is typically positioned directly upstream of the Bub operon, similar to the positioning of *capP-capH*, *capW*, and *capK-capS* (**Table S1; Figure 1A**). The first protein in this operon is predicted to contain a σ^4^ DNA-binding domain (Pfam PF04545) found in bacterial sigma factors; the second protein contains no predicted domains; and the large third protein contains a predicted HExxH zinc metallopeptidase domain (PF06114), a UvrD-like helicase domain (PF13361), and a PD-(D/E)xK family nuclease domain (PF12705) (**Figure 1A**). The presence of a predicted sigma factor, plus this operon’s consistent location upstream of Bub operons, suggested that these three proteins function together as a transcriptional regulator system. The presence of three distinct enzymatic domains in the large third protein led us to term this system “Cerberus” after the three-headed “Hound of Hades” that guards the entrance to the underworld in Greek mythology, with individual genes denoted *cerA* (predicted sigma factor), *cerB* (uncharacterized), and *cerC* (predicted peptidase-helicase-nuclease). We performed comprehensive BLAST searches with *cerC* in the Integrated Microbial Genomes (IMG) database, and identified 520 Cerberus operons (**Table S2**). In 500 cases, Cerberus is encoded directly upstream of a verified or predicted defense/immune pathway, including type II CBASS (cyclic oligonucleotide-based anti-phage signaling system; 14 cases)^23,24^, type I/II Bil (bacterial ISG15-like; 17 cases)^8,25,26^, and type I/II Bub (469 cases). We also identified nine cases where Cerberus is encoded upstream of an operon encoding homologs of the well-characterized DNA repair proteins SbcC and SbcD (**Figure 1B**)^27^.

**Figure 1.**
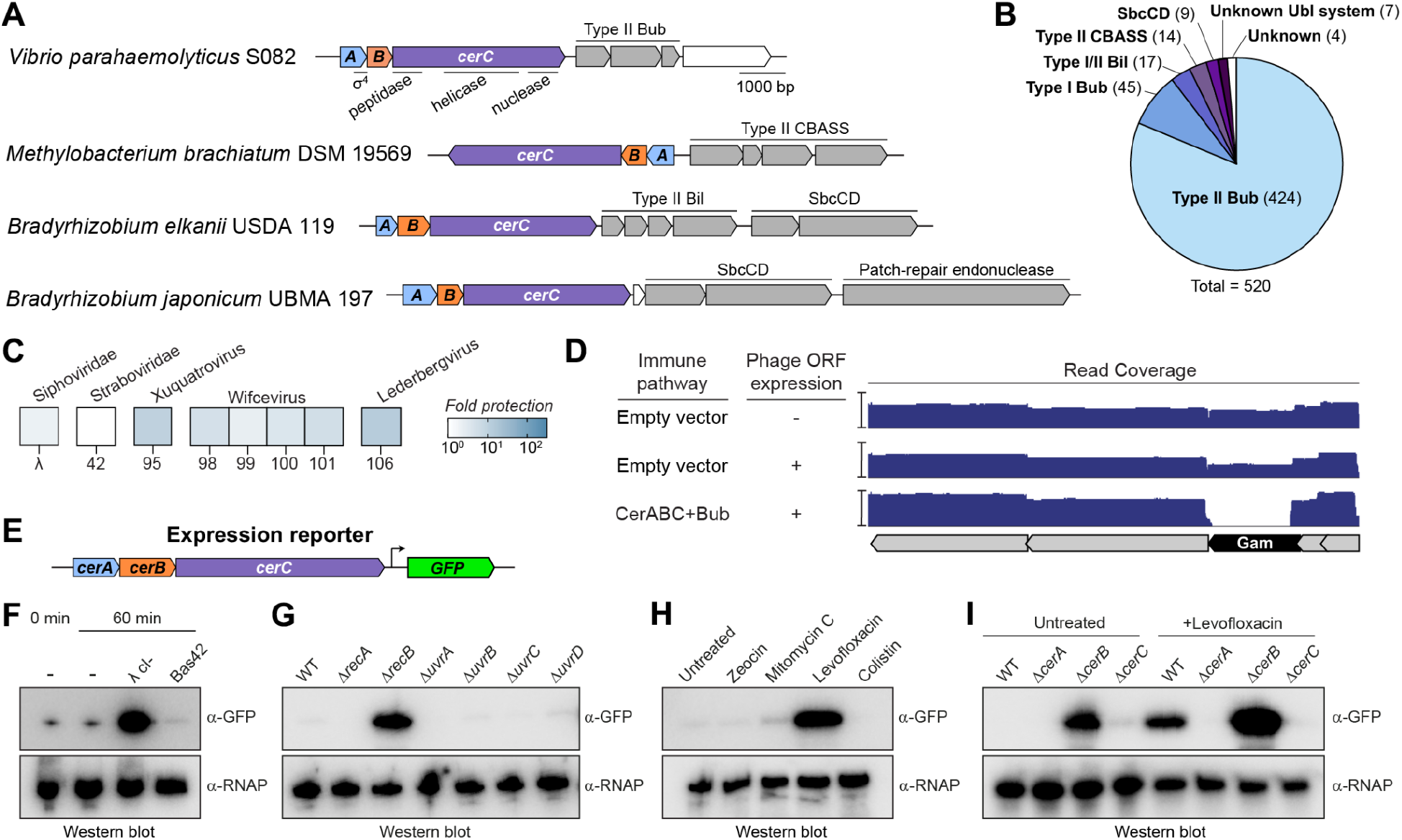
Cerberus is associated with known and putative defense/immune operons. A. Four Cerberus operons comprising *cerA* (light blue), *cerB* (orange), and *cerC* (purple) and their neighboring operons. For one operon from *V. parahaemolyticus* (top), automatically annotated domains for CerA (σ^4^, PF04545) and CerC (peptidase, PF06114; helicase, PF13361; nuclease, PF12705) are noted below each gene. Genes colored white are unknown. See **Table S2** for a full list of identified Cerberus operons. B. Pie chart showing identified operons neighboring Cerberus operons. See **Table S2** for details. C. Heat map of immunity provided by the *V. parahaemolyticus* Cerberus-Bub combined operons in an *E. coli* host. See results from full screen in **Figure S1**. D. Sequencing read depth across a region of the phage λ genome in different conditions, indicating that expression of λ Gam is toxic in the presence of the *V. parahaemolyticus* Cerberus-Bub combined operons. See **Figure S2** for read coverage across the full λ genome. E. Schematic of the GFP expression reporter used for panels F-I, with the *V. parahaemolyticus* Cerberus operon and native downstream promoter, with the Bub operon replaced with a gene encoding GFP. F. Expression reporter showing GFP expression 60 minutes after infection with either phage λ (*cI*^-^mutant) or Bas42. Top panel shows an anti-GFP western blot, and bottom panel shows an anti-RpoB (RNA polymerase subunit) western blot as loading control. G. Expression reporter showing GFP expression in *E. coli* Keio strains knocked out for the indicated DNA repair genes. See **Materials and Methods** for strain IDs. H. Expression reporter showing GFP expression in the presence of selected compounds. Zeocin, mitomycin C, and levofloxacin cause DNA damage through different mechanisms, and colistin is a membrane-disrupting drug. I. Expression reporter showing GFP expression in the presence or absence of levofloxacin, with the expression reporter vector knocked out for *cerA*, *cerB*, or *cerC*.

### Cerberus regulates transcription of its associated operon

The association of Cerberus with predicted anti-phage immune pathways suggests that it may be activated by phage infection. We tested the ability of a Cerberus-type II Bub locus from *Vibrio parahaemolyticus* S082 to protect an *E. coli* host against infection by phages in the extended BASEL collection^28^. The Cerberus-Bub locus provided modest protection (up to 2 logs in a plaque assay) against a range of phages, including members of *Siphoviridae* (notably phage λ_vir_), the subfamily *Guernseyvirinae*, and the genera *Skatevirus* and *Wifcevirus* (**Figure 1C, Figure S1**). We previously developed an assay to identify phage-encoded triggers for immune pathways that measures depletion of specific phage ORFs from a plasmid library upon co-transformation with a cytotoxic immune pathway^29^. Reasoning that the *V. parahaemolyticus* Cerberus-Bub locus likely inhibits cell growth upon sensing phage infection^22^, we used this assay to search for Cerberus-activating triggers in the phage λ proteome. We found that in the presence of the *V. parahaemolyticus* Cerberus-Bub locus, plasmids encoding the RecBCD inhibitor Gam were specifically depleted, indicating that λ Gam activates cytotoxicity by Cerberus-Bub (**Figure 1D, Figure S2**).

To directly test whether the Cerberus operon regulates expression of its associated immune/defense operon, we constructed a GFP expression reporter by replacing the downstream type II Bub operon from *V. parahaemolyticus* S082 with a gene encoding GFP, retaining the Cerberus operon and the intergenic sequence downstream of *cerC* (**Figure 1E**). *E. coli* cells carrying the *V. parahaemolyticus* Cerberus GFP reporter showed high GFP expression upon infection with phage λ (**Figure 1F**). Expression of λ Gam alone, in the absence of infection, was also sufficient to trigger GFP expression (**Figure S3**). Conversely, infection with the T4-like phage Bas42, against which the *V. parahaemolyticus* Cerberus-Bub locus did not provide protection, did not activate GFP expression (**Figure 1F**). To confirm a specific role for RecBCD inhibition in Cerberus-mediated expression, we transformed the GFP reporter into *E. coli* Keio strains^30^ lacking individual DNA repair proteins, including *recA*, *recB*, *uvrA, uvrB, uvrC,* or *uvrD*. In this experiment, we observed GFP expression only in *recB* knockout cells (**Figure 1G**). Together, these results indicate that Cerberus mediates expression of its associated immune/defense operon, in response to the inhibition or loss of RecBCD. Finally, we tested whether high levels of DNA damage (which could overwhelm RecBCD’s repair capacity) could also activate Cerberus, and found that addition of the type II DNA topoisomerase inhibitor levofloxacin triggered GFP reporter expression (**Figure 1H**).

To determine the role of each gene—*cerA*, *cerB*, and *cerC*—in Cerberus-mediated expression, we generated individual gene deletions in our GFP expression reporter, and tested for GFP expression in the presence and absence of levofloxacin (**Figure 1I**). We found that deletion of either *cerA* or *cerC* eliminated GFP expression in the presence of levofloxacin. Conversely, deletion of *cerB* resulted in high-level GFP expression even in the absence of levofloxacin, indicating that CerB is a negative regulator of Cerberus-mediated expression.

### CerB is an anti-sigma factor that negatively regulates CerA

Our GFP reporter assays showed that CerB acts as a negative regulator in the Cerberus system, suggesting that it may act as an “anti-sigma factor” by binding and inhibiting the activity of the predicted sigma factor CerA. Supporting this idea, we could co-purify a complex of *V. parahaemolyticus* CerA and CerB (**Figure 2A**). We used AlphaFold 3 to predict the structure of this complex, resulting in a model with an interface predicted template modeling score (ipTM) of 0.81, indicating high confidence that CerA and CerB form a complex (**Figure 2B, Figure S4**). The predicted structure shows CerA with an N-terminal σ^2^ domain and a C-terminal σ^4^ domain, an architecture characteristic of group IV sigma factors, including the *E. coli* alternative sigma factor σ^E^ ^31^. Group IV sigma factors are typically associated with anti-sigma factors that directly bind and inhibit their cognate sigma factors; protease-mediated destruction of the anti-sigma factor in response to extracellular or cytoplasmic signals results in sigma factor release and activation^32,33^. In keeping with this model, the predicted CerA-CerB structure shows CerB wrapping around CerA, forming an overall architecture similar to known structures of group IV sigma factors bound to their cognate anti-sigma factors (**Figure 2B-C**)^31^.

**Figure 2.**
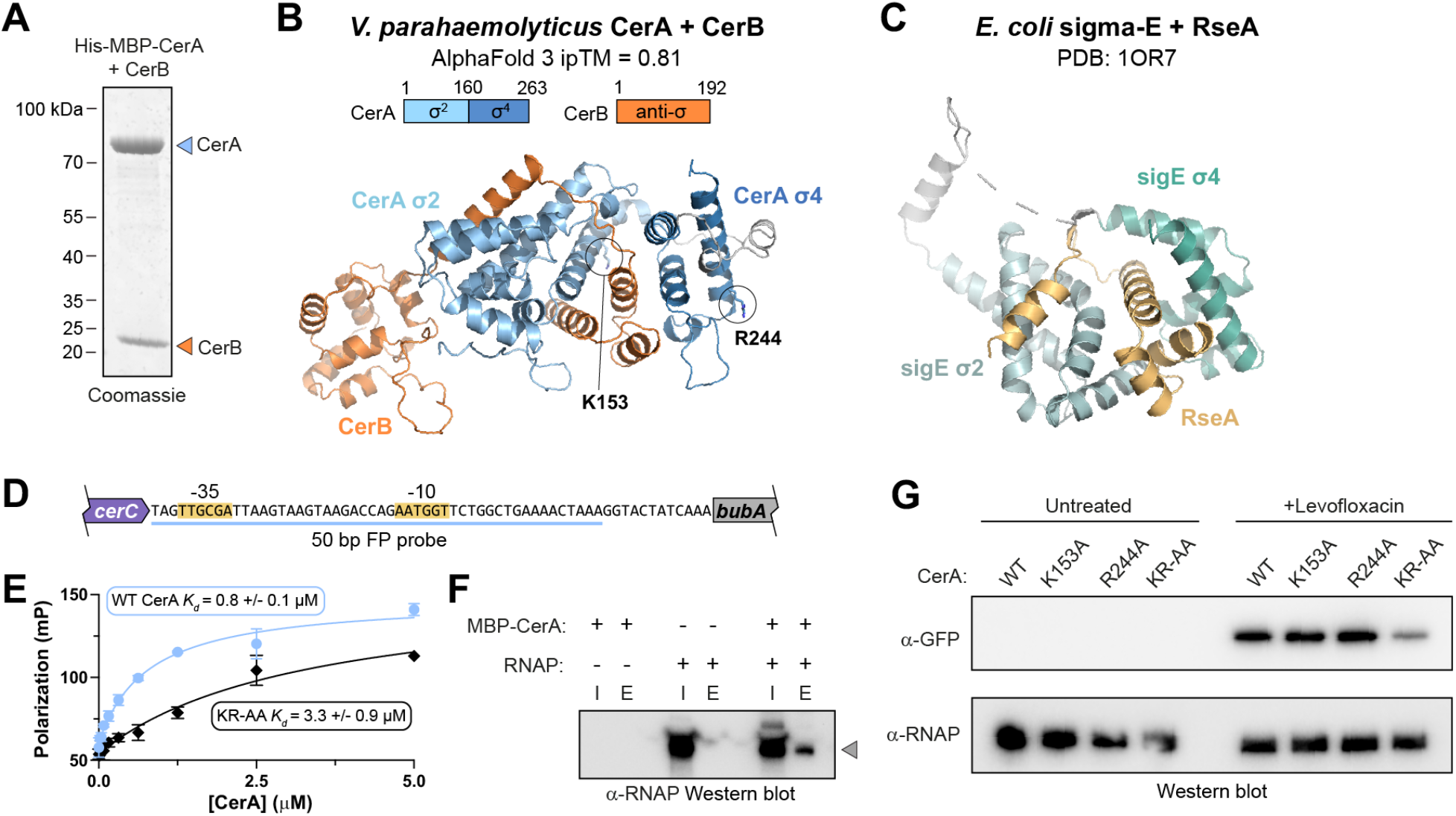
CerA and CerB constitute a sigma factor–anti-sigma factor pair. A. Coomassie-stained SDS-PAGE gel showing a purified complex of *V. parahaemolyticus* CerA (His_6_-MBP tagged) with untagged CerB. B. AlphaFold 3 model of *V. parahaemolyticus* CerA-CerB complex (see **Figure S4** for details). C. Structure of *E. coli* σ^E^ bound to its anti-sigma factor RseA (PDB 1OR7)^31^, roughly aligned to the predicted CerA-CerB structure in panel (B). D. Sequence of the intergenic region between *V. parahaemolyticus cerC* and *bubA*, with predicted -35 and -10 sites (from PromoterCalculator: https://www.denovodna.com/software/), and the 50 bp probe used for fluorescence polarization (FP) assays denoted with a light blue line. E. Fluorescence polarization assay showing binding of *V. parahaemolyticus* CerA to the 50-bp double-stranded FP probe shown in panel (D). Light blue circles: wild type (WT) CerA; black diamonds: K153A/R244A double mutant. *K_d_* values were calculated using a single-site binding model from triplicate data. See **Figure S6** for results with single mutants. F. Anti-RNA polymerase β subunit (RpoB) western blot showing results of an amylose pulldown assay with His_6_-MBP tagged *V. parahaemolyticus* CerA (bait) and purified *E. coli* RNA polymerase. I: input fraction; E: amylose-bound elution fraction. G. GFP reporter assay showing effects of CerA point mutations. KR-AA: K153A/R244A double mutant.

Bacterial sigma factors promote gene expression by binding specific promoters and then recruiting RNA polymerase for transcription. We used AlphaFold 3 to model a structure of a promoter-bound CerA-RNA polymerase holoenzyme, which revealed the likely DNA binding surfaces of CerA’s σ^2^ and σ^4^ domains (**Figure S5**). Based on this prediction, we generated point mutations in the CerA σ^2^ domain (K153A) and σ^4^ domain (R244A) that are predicted to disrupt DNA binding. We used a fluorescence polarization-based assay to measure binding of purified His_6_-MBP tagged *V. parahaemolyticus* CerA to a 50-bp DNA from the intergenic region downstream of *cerC*, containing the putative promoter for the downstream type II Bub operon (**Figure 2D-E**). We found that CerA bound this DNA, and each single mutant (K153A and R244A) had only a minor effect on binding (**Figure S6**). Combining the mutants had a stronger effect, with the K153A/R244A double mutant showing reduced DNA binding (**Figure 2E**). We next performed an amylose pulldown assay using MBP-tagged *V. parahaemolyticus* CerA and purified core *E. coli* RNA polymerase, and detected a direct interaction (**Figure 2F**). Finally, we tested the CerA DNA-binding mutants in our GFP expression reporter, and found that the K153A/R244A double-mutant strongly reduced GFP expression upon addition of levofloxacin (**Figure 2G**). Together, these data suggest that CerA and CerB function as a sigma factor–anti-sigma factor pair that regulates expression of its associated defense/immune operon.

### Structure of CerC

Since anti-sigma factors of group IV sigma factors are typically regulated by protease-mediated degradation^32^, and CerC contains a predicted zinc metallopeptidase domain, we reasoned that CerC may become activated to cleave CerB upon the inhibition of RecBCD and/or the introduction of DNA damage. To learn more about CerC’s structural mechanisms, we sought to determine a 3D structure of this protein. We were unable to purify full-length *V. parahaemolyticus* CerC, but we could purify a CerC protein encoded by an *E. coli* Cerberus operon associated with a type II Bub operon (**Figure S7A**). We found that *E. coli* CerC co-purifies with bound DNA, which could be removed with a high-salt wash during the initial Ni^2+^ affinity chromatography step (see **Materials and Methods**). Using these purified samples, we determined separate cryoelectron microscopy (cryo-EM) structures of both apo CerC (purified with high-salt wash) and DNA-bound CerC (purified without high-salt wash) (**Figure 3A-C, Figure S7, Figure S8, Table S3**).

**Figure 3.**
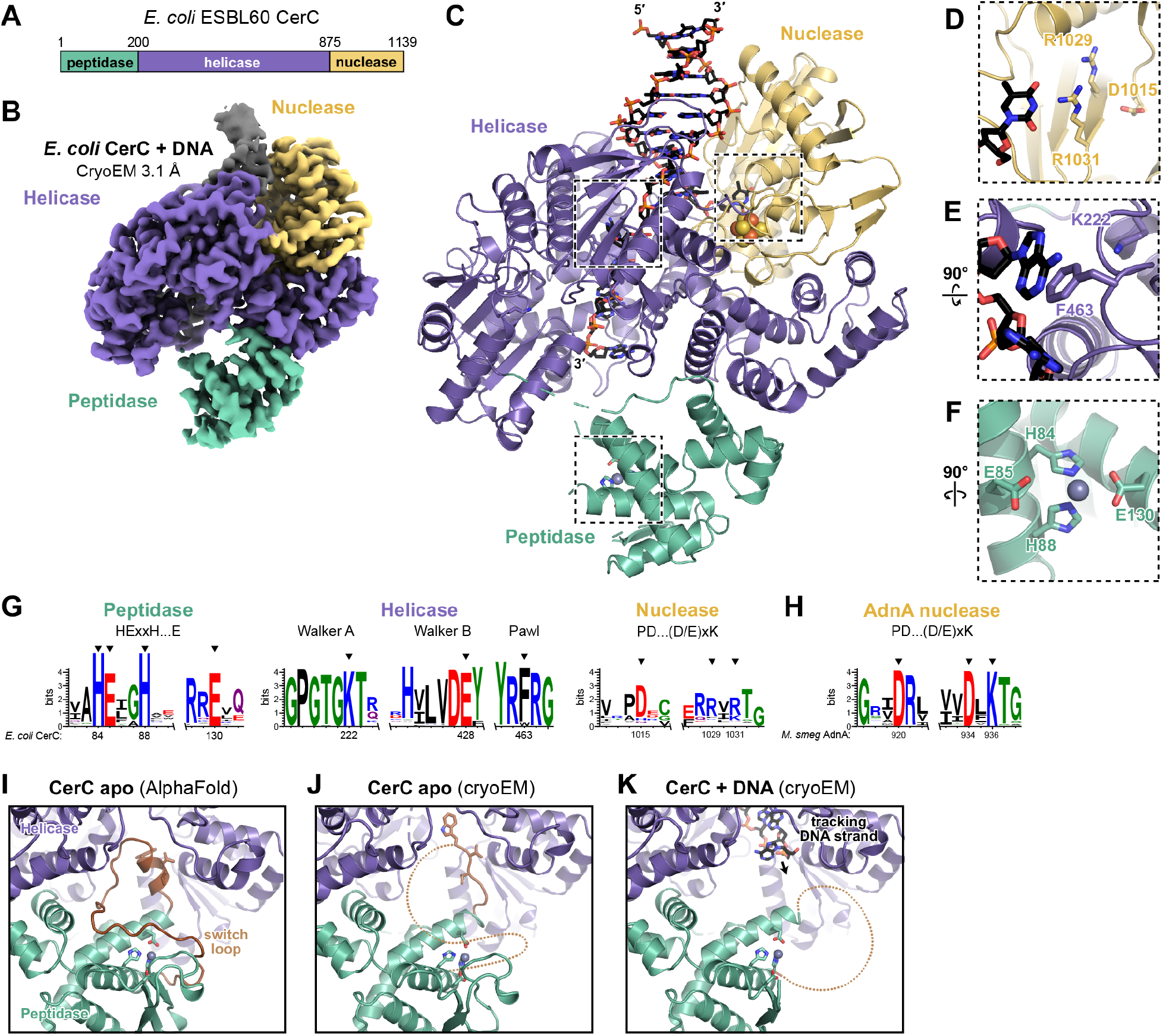
Structure of CerC. A. Domain diagram of *E. coli* ESBL60 CerC. B. Cryo-EM density for *E. coli* CerC bound to DNA. See **Figure S7** for apo CerC structure determination details, and **Figure S8** for CerC-DNA complex structure determination details. C. Overall structure of *E. coli* CerC bound to DNA, with peptidase domain green, helicase domain purple, and nuclease domain yellow. Bound DNA is shown as black sticks, and the 4Fe-4S cluster bound to the nuclease domain is shown as spheres. D. Closeup view of the nuclease active site, with residues corresponding to the canonical PD-(D/E)xK active site shown as sticks. One DNA strand is visible as sticks. E. Closeup view of the helicase active site and DNA-bound ratchet pawl. F. Closeup view of the peptidase active site with bound Zn^2+^ ion. G. Sequence logos for the CerC peptidase, helicase, and nuclease catalytic sites, built from 655 CerC sequences identified by PSI-BLAST^55^. Black arrowheads at top indicate critical functional residues. The residues corresponding to the PD-(D/E)xK nuclease catalytic site are not well conserved. H. Sequence logo showing high conservation of the catalytic residues in the active nuclease AdnA. I. Closeup of the putative switch loop from an AlphaFold 3 predicted structure of *E. coli* CerC (see **Figure S9** for details). The switch loop (shown in brown) is predicted to occlude the peptidase active site and dock against the inner surface of the DNA exit tunnel on the helicase domain. J. View as in panel (I) of the apo CerC structure, showing the partially ordered switch loop with hydrophobic residues docked against the DNA exit tunnel. K. View as in panel (I) of the CerC-DNA structure, showing that the switch loop is disordered and the tracking DNA strand is occupying the same space in the DNA exit tunnel.

Our structures reveal CerC as a monomeric protein with an N-terminal zinc metallopeptidase domain, central UvrD-type helicase domain, and C-terminal PD-(D/E)xK nuclease domain (**Figure 3C-F**). The N-terminal peptidase domain closely resembles the HExxH zinc metallopeptidases IrrE and CapP^15,34^. Our structures reveal a Zn^2+^ ion coordinated by conserved residues His84, His88, and Glu130 in the active site, adjacent to the putative catalytic residue, Glu85 (**Figure 3F-G**). CerC’s helicase and nuclease domains are tightly associated with one another, forming an overall architecture that closely resembles bacterial DNA repair-associated helicase-nuclease complexes RecBCD, AddAB, and AdnAB (see detailed discussion below)^35–37^. CerC’s helicase domain contains a highly conserved ATP-binding site with Walker A and B motifs, plus a conserved phenylalanine residue (Phe463) that in related helicase-nuclease enzymes acts as a “ratchet pawl” during DNA unwinding (**Figure 3E,G**)^38–40^. The nuclease domain contains a structural 4Fe-4S cluster coordinated by four conserved cysteine residues (Cys907, Cys1109, Cys1112, and Cys1118), similar to that found in the related helicase-nuclease proteins AdnA and AddB (see below).

Examination of the putative PD-(D/E)xK nuclease active site reveals poor conservation: while *E. coli* CerC possesses the canonical proline-aspartate pair (residues 1014-1015), these residues are not highly conserved across CerC proteins. CerC proteins uniformly lack the downstream aspartate/glutamate and lysine residues that are required for nuclease activity in this enzyme family, with both residues replaced with arginine in *E. coli* CerC (**Figure 3D,G,H**). The lack of key active site residues means that while the CerC C-terminal domain shares a 3D fold with PD-(D/E)xK nucleases, it is unlikely to be an active nuclease.

The structure of DNA-bound CerC captures the enzyme in the act of unwinding DNA. Because our DNA-bound CerC structure contains DNA that co-purifies with the enzyme from *E. coli*, and therefore represents a range of sequences, we modeled one strand of the DNA as poly-T and the other as poly-A. We observe a short segment of double-stranded DNA bound to the helicase and nuclease domains, with the two strands separating and the 3ʹ end of one strand (termed the tracking strand) passing through the helicase to emerge near the peptidase domain (**Figure 3C**). The conserved “ratchet pawl” residue Phe463 forms pi-stacking interactions with one base in the tracking strand (**Figure 3E**). The displaced 5ʹ strand extends into the nuclease domain, with ordered density for this strand ending near the diverged catalytic site (**Figure 3C-D**).

In previous work, we showed that the IrrE-like zinc metallopeptidase CapP contains a “switch loop” that occludes the peptidase active site, and is likely displaced upon ssDNA binding to activate the enzyme^15^. The related peptidase IrrE does not possess a switch loop, but is similarly activated by binding ssDNA^41^. An AlphaFold 3 model of CerC (**Figure S9**) reveals a putative switch loop similar to that in CapP (residues 93-124 of *E. coli* CerC) that is predicted to extend over the peptidase active site and dock against the helicase domain, at the terminus of the tunnel through which the tracking strand is extruded (**Figure 3I**). In our structure of CerC in the apo state, this loop is partially ordered, with several hydrophobic residues near the C-terminus of the loop (Trp119, Val120, and Val121) docked against the inner surface of the DNA tunnel in the helicase domain (**Figure 3J**). In our structure of DNA-bound CerC, no density for these residues is observed, and the tracking DNA strand emerging from the helicase domain occupies the same space (**Figure 3K**). Together, these observations suggest that DNA binding and unwinding by the CerC helicase and nuclease domains may regulate the protein’s peptidase activity by mediating conformational changes within this putative switch loop.

About one-third of CapP proteins possess a critical cysteine residue in their switch loop—termed the “cysteine switch”—that directly coordinates the catalytic Zn^2+^ ion to maintain the protein in an inactive state in the absence of ssDNA^15^. Neither *E. coli* nor *V. parahaemolyticus* CerC contain a cysteine residue on their switch loops, but in sequence alignments, we identify a putative cysteine switch residue in 32% of CerC proteins (**Figure S10**). This observation further supports the idea of a switch loop mechanism in CerC, with displacement of this loop by ssDNA emerging from the helicase domain triggering peptidase activation.

### CerC’s peptidase activity is controlled by DNA processing via its helicase and nuclease domains

Our structures of CerC suggest a mechanism by which CerC binds and unwinds DNA, with the resulting ssDNA activating the enzyme’s peptidase domain. To test the requirement for CerC’s three domains, we generated a series of point-mutations in the CerC peptidase, helicase (Walker A and ratchet pawl), and nuclease domains (**Figure 4A**). We first tested these mutations in our GFP expression reporter, and found that mutating the peptidase, helicase Walker A, and helicase ratchet pawl all eliminated GFP expression (**Figure 4B**). Mutating the poorly conserved aspartate in the nuclease domain strongly reduced, but did not eliminate, GFP expression (**Figure 4B**). Since the nuclease domain is unlikely to be catalytically active, we hypothesize that the nuclease-domain mutation reduces the DNA unwinding activity of the enzyme.

**Figure 4.**
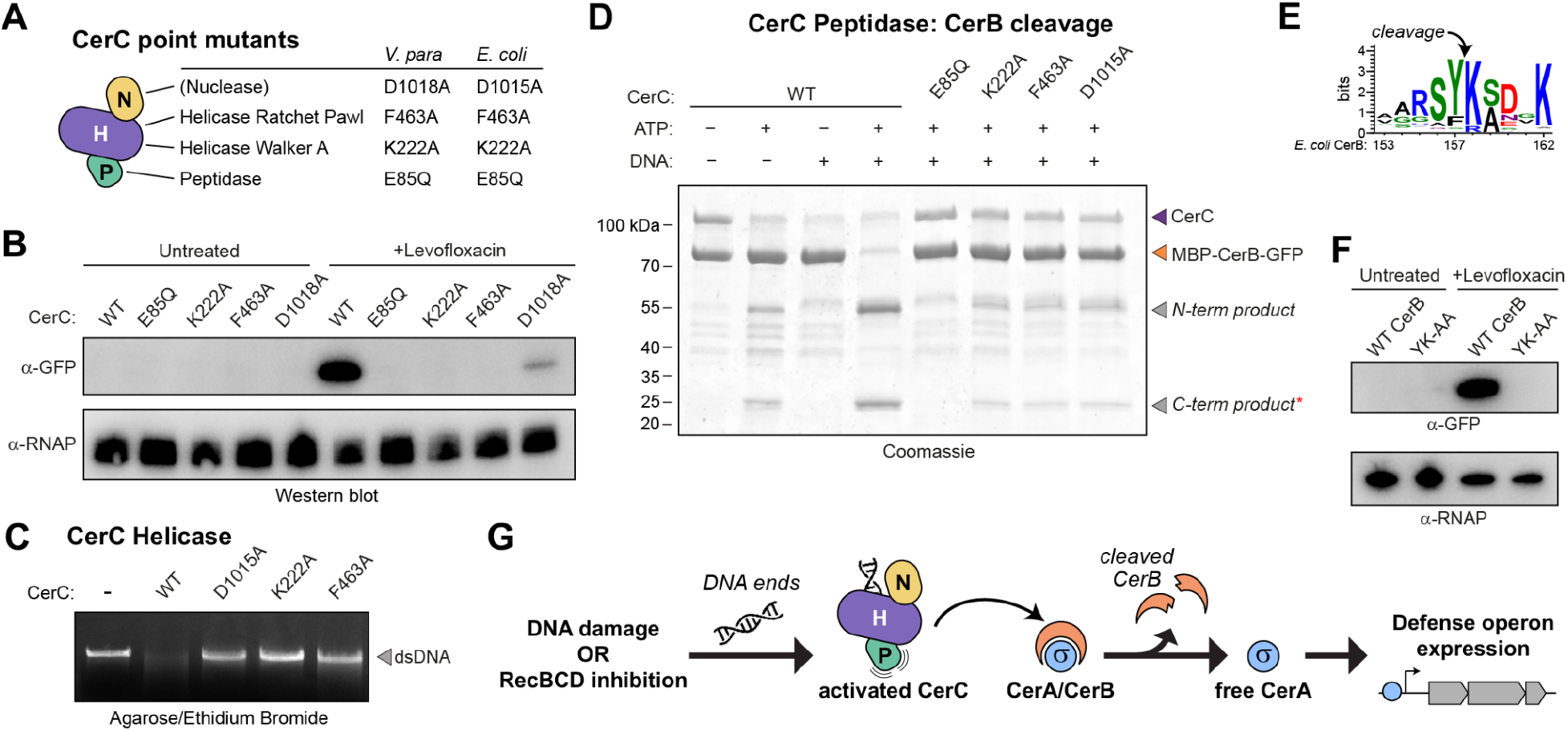
CerC is a CerB-specific peptidase activated by DNA binding and unwinding. A. Summary of point mutations in *V. parahaemolyticus* and *E. coli* CerC. In the cartoon schematic, the peptidase domain (P) is green, the helicase domain (H) is purple, and the nuclease domain (N) is yellow. The notation “(nuclease)” is intended to convey that CerC’s nuclease domain is likely to be catalytically inactive. B. GFP expression reporter assay for *V. parahaemolyticus* CerC mutants, in the absence and presence of levofloxacin. C. Helicase assay showing disappearance of an 11-kb linear double-stranded DNA when incubated with wild type (WT) *E. coli* CerC in the presence of ATP, and the disruption of this activity by mutations in the helicase and nuclease domains. D. *In vitro* CerB cleavage assay, using an *E. coli* MBP-CerB-GFP fusion substrate (orange arrowhead), WT or mutant CerC, DNA, and ATP. The C-terminal product of cleavage (asterisk) was isolated for Edman sequencing (panel E, **Figure S11**). E. Sequence logo of CerB, showing high conservation of the CerC cleavage site identified by Edman sequencing (see **Figure S11**). F. GFP expression reporter assay showing the effect of mutating the CerC cleavage site on CerB (Tyr157 and Lys158 mutation to alanine: YK-AA). G. Model for Cerberus function. DNA damage and/or inhibition of RecBCD leads to the buildup of DNA ends. Binding and unwinding of DNA by CerC activates its peptidase domain, triggering cleavage of CerB, release of CerA, and recruitment of RNA polymerase to boost expression of the Cerberus-associated defense/immune operon.

Next, we used purified *E. coli* CerC and tested helicase activity by incubating the enzyme with ATP and linearized plasmid DNA. While double-stranded DNA forms a compact band on an agarose gel that is readily stained with ethidium bromide, ssDNA forms a more diffuse band that does not readily stain with ethidium bromide. We found that wild type CerC readily eliminated the double-stranded DNA band in this assay, and that mutations in both the helicase and nuclease domains reduced this activity (**Figure 4C**).

We next incubated purified *E. coli* CerC with its cognate CerB protein, and found that CerB was robustly cleaved in the presence of both ATP and linear DNA (**Figure 4D**). Mutation of CerC’s peptidase catalytic glutamate (E85Q) eliminated CerB cleavage, while mutations in the helicase and nuclease domains strongly reduced cleavage (**Figure 4D**). We isolated the C-terminal cleavage fragment of CerB and subjected it to Edman degradation to determine the cleavage site, and found that cleavage occurs between Tyr157 and Lys158, which are highly conserved in CerB (**Figure 4E, Figure S11**). Mutating the equivalent residues in *V. parahaemolyticus* CerB (also Tyr157 and Lys158) to alanine in our GFP reporter eliminated GFP expression in the presence of levofloxacin, supporting the idea that CerB cleavage is required for Cerberus-mediated expression (**Figure 4F**).

We next probed the role of CerC’s peptidase-domain switch loop in CerB cleavage. We first purified the isolated *E. coli* CerC peptidase domain and a mutant lacking residues 93-124 (Δloop). We tested activity of this construct in the presence and absence of a 22-nt ssDNA oligonucleotide to mimic the single-stranded DNA generated by the CerC helicase domain. We found that the wild type peptidase domain was active in the absence of DNA, and was not further activated by ssDNA (**Figure S12A**). This observation suggests that in the absence of the CerC helicase domain, the switch loop is unable to inhibit CerC’s peptidase activity. Curiously, the Δloop mutant was markedly less active than the wild type peptidase domain, and was not stimulated by ssDNA (**Figure S12A**). This result suggests that the switch loop may play a positive role in substrate recognition, once undocked from the helicase domain. Supporting this idea, deleting the switch loop in full-length CerC completely eliminated CerB cleavage *in vitro* (**Figure S12B**). Together, these data suggest that CerC’s switch loop both inhibits the peptidase activity in the absence of DNA, and aids substrate recognition and cleavage once CerC is activated. AlphaFold 3 structure predictions of a CerC-CerB complex (in the presence or absence of DNA) failed to generate confident models, leaving the structural basis for CerB recognition unknown.

Overall, our combined cellular, biochemical, and structural data support a model for Cerberus activation that begins with RecBCD inhibition and/or DNA damage, generating free DNA ends that CerC binds and unwinds. The unwound tracking strand emerging from CerC’s helicase domain activates its peptidase domain, enabling CerB cleavage and the release of CerA to promote expression of the associated immune/defense operon (**Figure 4G**).

### CerC resembles bacterial helicase-nuclease DNA repair complexes

Our structures of CerC reveal strong structural similarity to bacterial DNA repair-associated helicase-nuclease complexes, including AdnAB, AddAB, and RecBCD (**Figure 5A-D**). Each of these complexes contains two fused helicase-nuclease proteins that work together to resect double-stranded DNA breaks as a first step in their repair. AdnAB is the simplest of these machines and is considered to be ancestral to AddAB and RecBCD^42^. Each protein in AdnAB, AddAB, and RecBC (RecD is a helicase-only protein) encodes helicase-like and nuclease-like domains, but these complexes differ in which of these domains retain catalytic activity (**Figure 5B-D**).

**Figure 5.**
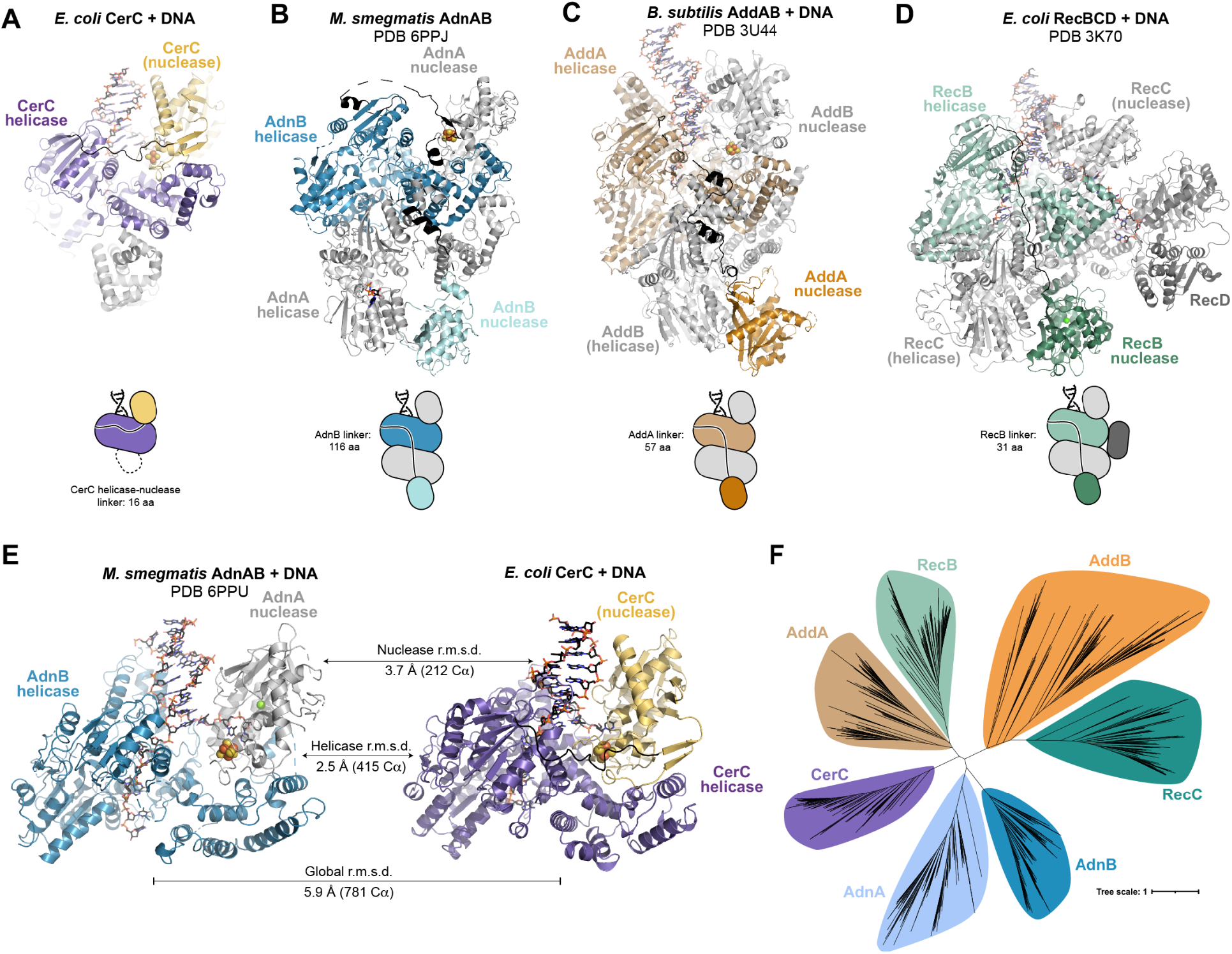
CerC is related to bacterial DNA-repair associated helicase-nuclease complexes. A. Overall structure of DNA-bound *E. coli* CerC, with peptidase domain gray, helicase domain purple, and nuclease domain yellow. The helicase-nuclease linker is shown in black. In panels (A)-(D), a domain name in parentheses, e.g., “(nuclease)” is intended to convey that the domain is catalytically inactive. B. Overall structure of *M. smegmatis* AdnAB (PDB 6PPJ)^38^, with AdnB colored blue and AdnA colored gray. C. Overall structure of DNA-bound *B. subtilis* AddAB (PDB 3U44)^75^, with AddA colored brown and AddB colored gray. D. Overall structure of DNA-bound *E. coli* RecBCD (PDB 3K70)^76^, with RecB colored green, RecC colored light gray, and RecD colored dark gray. E. Aligned structures of DNA-bound *M. smegmatis* AdnAB (left; PDB 6PPU)^38^ and *E. coli* CerC, showing the structural similarity between these complexes’ leading helicase-nuclease units, and the structural similarity between their component domains. F. Unrooted phylogenetic tree constructed from ∼150 examples of each bacterial helicase-nuclease protein family. CerC is not unambiguously more related to any specific helicase-nuclease family. See **Figure S13** for bootstrap values.

While their overall architecture varies, all three helicase-nuclease complexes engage substrate DNA with a “leading” helicase motor domain (from AdnB, AddA, and RecB, respectively) that translocates the tracking strand in a 3ʹ-to-5ʹ direction, and feeds the opposite strand into a tightly associated nuclease domain (from AdnA, AddB, and RecC, respectively) for 5ʹ-to-3ʹ degradation (**Figure 5A-D**; RecC’s nuclease domain is inactive, and this strand is instead degraded by RecD). The juxtaposition of CerC’s helicase and nuclease domains closely resembles these “leading helicase+nuclease” units, with the key difference being that the helicase and nuclease domains of CerC are in the same polypeptide, while the structurally equivalent helicase and nuclease domains of AdnAB, AddAB, and RecBC are in different polypeptides. Pairwise structural comparisons using DALI^43^ of CerC’s helicase and nuclease domains with AdnA/AdnB, AddA/AddB, and RecB/RecC show that CerC’s helicase domain most closely resembles the helicase domain of AdnB (DALI Z-score 25.5; **Table S4**), and that CerC’s nuclease domain most closely resembles the nuclease domain of AdnA (DALI Z-score 15.4; **Table S5**). AdnA, but not AdnB, also possesses a 4Fe-4S cluster in its nuclease domain similar to CerC (**Figure 5A-B**). Finally, CerC’s combined helicase and nuclease domains overlay well with the combined AdnB helicase + AdnA nuclease domains (**Figure 5E**).

Our structural data suggest that CerC may have arisen from an AdnAB-like complex in one of two ways. First, fusion of an AdnA/AdnB-like protein to an N-terminal peptidase domain could have eliminated the protein’s ability to dimerize with its partner, forcing the helicase and nuclease domains to associate within the same polypeptide. Alternatively, a deletion within a tandem *adnB*-*adnA* operon that fused AdnB’s helicase domain to AdnA’s nuclease domain could have given rise to the single-polypeptide CerC assembly, followed by later fusion to a peptidase domain and inactivation of the nuclease domain. This scheme would also explain the presence of an aromatic ratchet pawl residue in CerC’s helicase domain (shared with AdnB) and a 4Fe-4S cluster in its nuclease domain (shared with AdnA). To gather support for an inferred evolutionary trajectory for CerC, we performed phylogenetic analysis of CerC’s helicase domain against the helicase domains of other bacterial helicase-nuclease proteins; unfortunately, this analysis was unable to define a convincing lineage for CerC (**Figure 5F, Figure S13**).

### Orthrus operons are Cerberus-like operons with a CapP-like peptidase

In our sequence-based searches for Cerberus operons, we noticed instances of a three-gene operon with a CerA-like predicted sigma factor, a CerB-like predicted anti-sigma factor, and a CapP-like zinc metallopeptidase. We named these operons Orthrus, after the two-headed sibling of Cerberus in Greek mythology, with individual genes denoted *ortA* (sigma factor), *ortB* (anti-sigma factor), and *ortC* (peptidase). Comprehensive BLAST searches in the IMG database identified 400 Orthrus operons, which are associated with a broad set of known or putative defense/immune operons including Pycsar, type II CBASS, and type I/II Bub (**Figure 6A-B, Table S6**). AlphaFold 3 modeling of an OrtA-OrtB complex from *Paraburkholderia graminis* predicts that these proteins form an intertwined sigma–anti-sigma complex (**Figure 6C, Figure S14A-B**). Modeling an OrtC-OrtB-ssDNA complex reveals the likely OrtC cleavage site on OrtB at a highly conserved Ala-Ala-Arg motif (**Figure 6D-F, Figure S14C-D**), and reveals a highly conserved cysteine on the OrtC switch loop that likely acts as a cysteine switch (**Figure 6G-H, Figure S14E-F**). These features of OrtC strongly resemble CapP (**Figure 6I-J**)^15^, and support a functional model for Orthrus that involves OrtC peptidase activation by ssDNA binding and cysteine switch disengagement, cleavage of OrtB, and release of OrtA to boost expression of an associated defense/immune operon.

**Figure 6.**
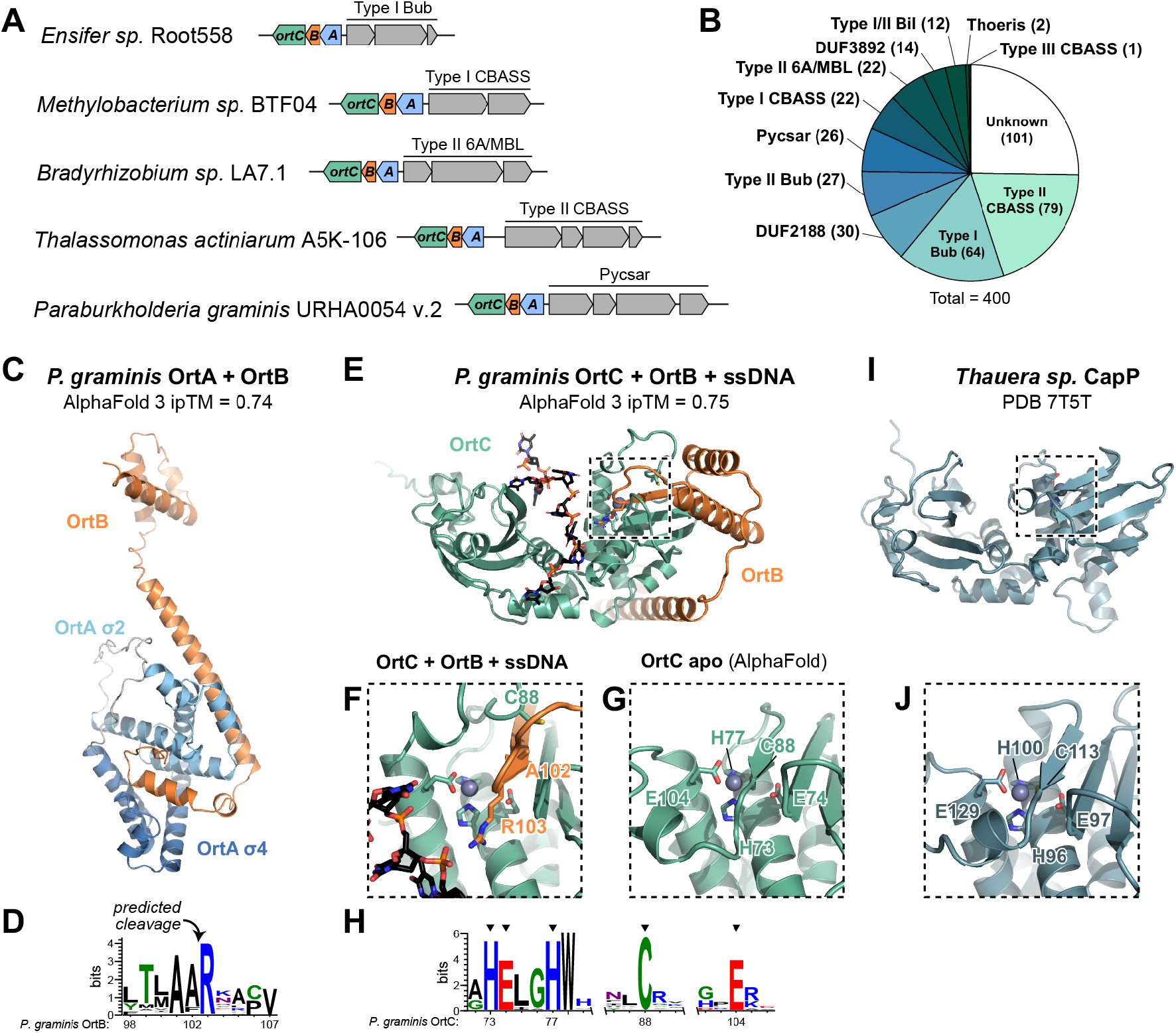
Orthrus operons resemble Cerberus and are associated with defense/immune operons. A. Five Orthrus operons comprising *ortA* (light blue), *ortB* (orange), and *ortC* (green) and their neighboring operons. See **Table S6** for a full list of identified Orthrus operons. B. Pie chart showing identified operons neighboring Orthrus operons. See **Table S6** for details. C. AlphaFold 3 model of an OrtA-OrtB complex from *Paraburkholderia graminis*, with OrtA colored blue (σ^2^ and σ^4^ domains labeled) and OrtB orange. See **Figure S14A-B** for AlphaFold 3 confidence metrics. D. Sequence logo of the predicted OrtC cleavage site on OrtB, based on sequence conservation and AlphaFold 3 modeling. E. Alphafold 3 model of a *P. graminis* OrtC-OrtB-ssDNA complex, with OrtC green, OrtB orange, and ssDNA black and shown as sticks. See **Figure S14C-D** for AlphaFold 3 confidence metrics. F. Closeup view of the OrtC active site in the AlphaFold 3 model of an OrtC-OrtB-ssDNA complex, with predicted cleavage-site residues on OrtB labeled. G. Closeup view of the OrtC active site in an AlphaFold 3 model of OrtC, with predicted active-site residues labeled. See **Figure S14E-F** for full model and AlphaFold 3 confidence metrics. H. Sequence logo of OrtC predicted active-site residues, including the HExxH…E motif and the cysteine switch. I. Crystal structure of *Thauera* sp. CapP (PDB 7T5T)^15^, oriented equivalently to OrtC in panel (E). J. Closeup view of the CapP active site, with active-site residues labeled.

## DISCUSSION

Here we identify Cerberus, a three-protein system that regulates expression of an associated defense/immune pathway in response to DNA damage and/or inhibition of the central DNA repair and stress-signaling complex RecBCD. Cerberus joins CapW/BrxR^16–18,20^, CapP-CapH^15^, and CapK-CapS^19^ as DNA damage-responsive transcriptional regulator systems associated with a variety of defense/immune operons including CBASS, bacterial ubiquitin-related pathways (Bil, Bub), and many others. Cerberus is sometimes associated with operons encoding homologs of the SbcCD DNA repair complex, suggesting that it plays a role in DNA repair in addition to anti-phage immunity. In contrast to the other regulators we identified, Cerberus does not depend on upstream DNA repair factors to generate ssDNA byproducts to activate signaling. Rather, CerC binds directly to broken DNA ends, enabling it to respond even when DNA repair machinery (notably RecBCD) is inhibited by phage-encoded factors.

CerC shares an overall architecture with bacterial DNA repair-associated helicase-nuclease complexes, including AdnAB, AddAB, and RecBCD. Specifically, CerC’s helicase and nuclease domains are equivalent to these complexes’ leading helicase-nuclease units. In AdnAB, AddAB, and RecBCD, these units are assembled from the helicase domain of one protein (e.g., AdnB) and the nuclease domain of its binding partner (e.g., AdnA). While sequence alignments do not reveal an unambiguous evolutionary path for CerC, structure-similarity searches show that its helicase domain most closely resembles AdnB, and its nuclease domain most closely resembles AdnA. Based on these observations, we propose that CerC evolved from an AdnAB-like complex, followed by incorporation of an N-terminal peptidase domain and loss of catalytic activity in its nuclease domain.

HExxH family zinc metallopeptidases play key regulatory roles in genetic conflicts of bacteria, phages, and mobile elements. In the most common paradigm, HExxH peptidases and their cognate transcriptional repressor targets control transcriptional switches that mediate mobilization of prophages (e.g., *Bacillus subtilis* prophage phi105) and integrative and conjugative elements (ICE elements, e.g., *Bacillus subtilis* ICE*Bs1*)^44,45^, activate general DNA damage responses (e.g., *Deinococcus* IrrE-DdrO)^34,46^, and drive expression of a range of immune pathways (e.g., CapP-CapH). HExxH peptidases are also paired with other cognate target proteins to control different signaling mechanisms. For example, the Orthrus pathways we identify here involve a peptidase (OrtC) that likely cleaves an anti-sigma factor (OrtB) to release a sigma factor (OrtA), thereby driving expression of an associated defense/immune operon. Recent work on the *M. tuberculosis* VapBC52 system revealed that the antitoxin VapB52 undergoes self-cleavage upon binding ssDNA, thereby relieving inhibition of the toxin VapC52 ^47^. CerC is the most diverged example to date, with the peptidase regulated by movement of its switch loop in response to DNA unwinding by a fused helicase domain, rather than by direct ssDNA binding. Nonetheless, all of these pathways share key features, notably activation by DNA damage and target cleavage at a similar consensus motif, typically between a bulky hydrophobic residue and a positively charged residue (CerC: Tyr-Lys; CapP: Phe-Arg; IrrE: Leu-Arg; VapB52: Phe-Arg [predicted]; OrtC: Ala-Ala-Arg [predicted])^15,46,47^.

The vast majority of identified Cerberus operons are associated with Bub pathways, the biological roles of which remain unclear. We previously showed that an *E. coli* Bub operon conjugates a ubiquitin-like protein to two proteins that participate in translation: the ribosomal protein RpsQ and the mRNA chaperone CspA^22^. Tellingly, 72% of Bub operons are associated with DNA damage-sensing transcriptional regulators, including CapW, CapP-CapH, CapK-CapS, and Cerberus (**Table S1**). We find that an additional ∼10% of Bub operons are associated with predicted transcription factors, or with toxin-antitoxin systems known to regulate their own expression in response to stress^48,49^. These systems could regulate their adjacent Bub operon by acting on their shared promoter region. Thus, around 82% of Bub operons are associated with known or putative stress-induced transcriptional regulators. Based on these observations, we propose that Bub pathways may not be dedicated anti-phage immune pathways; rather, they may slow metabolism (likely by inhibiting translation) in response to DNA damage or other stress. This response could manifest as immunity against phages that cause widespread DNA damage and/or inhibit RecBCD, but could also provide a survival advantage in other conditions aside from phage infection.

Both Cerberus and Orthrus are associated with individual genes and operons with unknown functions, highlighting the possibility of discovering new defense/immune pathways by their association with stress-sensing transcriptional regulators. Many of these pathways likely respond to threats other than phage infection, notably including DNA damage arising from other sources. This idea highlights the concept that defense islands are enriched not just for anti-phage immune pathways, but also for pathways that respond to other stresses. Future efforts to discover and mechanistically dissect these diverse pathways have the potential to unlock a more complete understanding of how bacteria sense and respond to diverse threats to their survival.

## MATERIALS AND METHODS

### Bioinformatics

Initial identification and annotation of *cerABC* genes associated with Bub operons^22^ was performed by manual inspection of gene neighborhoods using IMG (https://img.jgi.doe.gov) (**Table S1**). For comprehensive identification of *cerABC* associated immune operons, exhaustive BLAST searches in IMG were performed with representative CerC proteins, followed by manual annotation of genes encoding CerA and CerB (**Table S2**). Genomic DNA sequences ± 10 kb of each *cerABC* locus were downloaded from IMG and submitted to PADLOC^50^ for identification of nearby immune operons. Gene neighborhoods were further annotated by manual inspection in IMG.

For identification of *ortABC* operons, representative OrtC homologs were used for exhaustive BLAST searches in IMG and the associated *ortA* and *ortB* genes were manually annotated (**Table S6**). To identify nearby immune operons, ± 10 kb of each *ortABC* locus was downloaded and submitted to PADLOC. Gene neighborhoods were annotated by manual inspection in IMG.

To generate sequence logos of conserved residues in CerC and CerB, sequences of identified CerC and CerB proteins were aligned using the MAFFT version 7 web server (https://mafft.cbrc.jp/alignment/server/index.html)^51^ and visualized/filtered for redundancy in Jalview^52^. Sequence logos were generated with WebLogo version 3.9.1^53^.

To build a phylogenetic tree of the CerC, AddAB, RecBC, and AdnAB helicase domains, the accession numbers for all identified homologs of AddA, AddB, RecB, RecC, AdnA, and AdnB were obtained from the phylogenetic analysis performed in Mutte et al. (2025) and used as starting sequence sets^54^. For CerC, we used the NCBI BLAST webserver^55^ (NCBI BLAST: https://blast.ncbi.nlm.nih.gov/Blast.cgi) to identify 1000 candidate CerC homologs. Sequences shorter than 1000 aa and longer than 1300 aa were removed and the remaining sequences in each set were aligned using the MAFFT version 7 web server (https://mafft.cbrc.jp/alignment/server/index.html). Sequences with >95% redundancy were removed from the alignment using Jalview^52^, and a final experimental set of 150 sequences for each homolog was randomly selected with SeqKit version 2.13.0 ^56^ “sample2” operation. The remaining CerC sequences were analyzed with hmmscan in the HMMER webserver^57^ (https://www.ebi.ac.uk/Tools/hmmer/search/hmmscan) to confirm that all candidates encoded peptidase, helicase, and nuclease domains, and two sequences were removed for lacking N-terminal peptidase annotations.

Boundaries of the helicase domains for each homolog were identified using hmmscan in the HMMER webserver^57^. The experimental sequence set for each protein was aligned using the MAFFT version 7 web server and the helicase domains were extracted. The helicase domain sequences for all homologs were then aligned with MAFFT version 7.526 ^58,59^ using the “genafpair” (E-INS-i) algorithm with a maximum of 1000 iterations. The aligned columns were then trimmed with ClipKIT version 2.14.0 ^60^ using the “gappy” mode with a threshold of 0.9. The phylogenetic tree was built using IQ-TREE version 3.1.3 ^61^ with “Q.pfam+F+R10” as the model of evolution selected using the built in ModelFinder^62^, a maximum of 1000 Ultrafast bootstrap^63^ and 1000 replicates of the SH-like approximate likelihood ratio test (SH-aLRT)^64^. Trees were visualized in iTOL version 7.6 ^65^.

### Plaque assays

Double agar overlay assay was used to determine phage titer^66^. Briefly, overnight cultures of *E. coli* carrying plasmids expressing sfGFP (considered an empty vector control) under a low, constitutive promoter or the *V. parahaemolyticus* S082 *cerABC* + Bub locus under its native promoter in LB supplemented with carbenicillin (100 μg/mL) were diluted 1:100 into the same media and cultivated for 1.5 hours to reach logarithmic growth phase (A_600_ = 0.3 -0.8). 400 μL of the culture was then mixed with 3.5 mL of 0.35% agar LB with 10 mM MgCl_2_, 10 mM CaCl_2_, and 0.1 mM MnCl_2_ and poured over a 1.6% agar LB plate. After cooling at room temperature for 15 to 30 minutes, 2 μL of phage dilution series in SM Buffer was spotted onto the overlay and allowed to adsorb for about 10 minutes. The plates were then incubated overnight at 37°C and plaque forming units were enumerated the following day. For spots with hazy zones of clearances at the lowest concentration, compared to individual countable plaques, ten plaques were enumerated. For instances where no zone of clearance or plaque formation was observed at the least dilute spot, 0.9 plaques were enumerated as the limit of detection. Fold protection was calculated by dividing the phage titer enumerated on the empty vector strain by the phage titer enumerated on the defense system containing strain.

### Phage protein depletion screen

Identification of phage-encoded triggers of the *V. parahaemolyticus* S082 *cerABC* + Bub system were conducted as previously described^29^. Briefly, 50 μL of electrocompetent *E. coli* cells expressing GFP or the *V. parahaemolyticus* S082 *cerABC* + Bub genes under its native promoter were transformed with 180 ng of the phage λ ORFeome library. Cells were recovered at 37°C for 1 hour in S.O.C. with shaking at 220 rpm, serially-diluted and plated on MMCG agar plates (47.8 mM Na_2_HPO_4_, 22 mM KH_2_PO_4_, 18.7 mM NH_4_Cl, 8.6 mM NaCl, 22.2 mM Glucose, 2 mM MgSO_4_, 100 µM CaCl_2_, 3 µM Thiamine, Trace Metals at 0.1× ((Trace Metals Mixture T1001, Teknova; final concentration: 8.3 μM of FeCl3, 2.7 μM of CaCl_2_, 1.4 μM of MnCl_2_, 1.8 μM of ZnSO_4_, 370 nM of CoCl_2_, 250 nM of CuCl_2_, 350 nM of NiCl_2_, 240 nM of Na2MoO_4_, 200 nM of Na_2_SeO_4_ and 200 nM of H_3_BO_3_) plates supplemented with 100 μg/mL carbenicillin and 20 μg/mL chloramphenicol, with or without 50 μM IPTG. Plates were incubated at 37°C overnight and the plasmids from surviving colonies were isolated by miniprep. 20 μL of the miniprep was sequenced using 200Mbp Illumina Sequencing (SeqCenter), and processed as follows: Reads were mapped to the phage λ genome using the *Map to Reference* feature of Geneious Prime (default settings). Relative read abundance was compared using the Transcripts Per Million (TPM) feature in Geneious across three samples: (1) the empty vector and the phage λ ORFeome plated without IPTG, (2) the empty vector and the phage λ ORFeome plated with IPTG, and (3) the defense system and the phage λ ORFeome plated with IPTG. ORFs with 10 or less reads under non-inducing conditions were considered universally toxic and were unable to be assessed as activators of the system.

### GFP expression reporter assays

For GFP reporter expression assays, *E. coli* MG1655 cells were transformed with GFP expression reporter plasmids (wild type or indicated mutants), and 2 mL cultures in LB media plus 100 μg/mL ampicillin were grown overnight at 37°C with shaking. The next morning, 30 μL of saturated overnight culture was diluted into 3 mL fresh media and grown to an A_600_ of 0.3. Cells were stressed by the addition of 5 μg/mL mitomycin C, 5 μg/mL levofloxacin, 1 μg/mL colistin, or 50 μg/mL zeocin and grown for a further 1 h at 37°C. For GFP expression analysis, 500 μL was centrifuged, then the cell pellet was resuspended in 50 μL 2x SDS-PAGE loading buffer (125 mM Tris-HCl pH 6.8, 20% Glycerol, 4% SDS, 200 mM DTT, 0.01% w/v bromophenol blue). Sample volumes were adjusted based on the culture density to normalize the cells/volume in each sample. Samples were placed in a boiling water bath for 3 min, then 5 μL samples were loaded onto a TGX Stain-Free FastCast Acrylamide SDS-PAGE gel (Bio-Rad, 1610185) and separated at 180 V for 40 min at room temperature.

For GFP expression reporter assays in single-gene knockout strains, single-gene knockout *E. coli* strains from the Keio collection^30^ were obtained from Horizon Discovery (Strain IDs: parent strain BW25113^67^; *ΔrecA* JW2669, *ΔrecB* JW2788, *ΔuvrA* JW4019, *ΔuvrB* JW0762, *ΔuvrC* JW1898, *ΔuvrD* JW3786).

### Western blots

For western blots, samples were separated on 4-20% MiniPROTEAN TGX gels (Bio-Rad, 4561096), transferred to PVDF membranes (Bio-Rad Turbo transfer kit, 170-4272), and blocked for 1 h at room temperature with 5% nonfat dry milk in TBST (50 mM Tris-HCl pH 8.5, 150 mM NaCl, 0.1% Tween-20) followed by blotting for primary antibody at room temperature for 1 h (Antibodies used: Mouse anti-GFP primary antibody (Roche, 11814460001) at 1:3000 dilution; Mouse anti-RNA polymerase primary antibody (clone NT63; BioLegend #10019-878) at 1:3000 dilution). Blots were washed three times in TBST and incubated with Goat anti-mouse HRP-linked secondary antibody (Millipore Sigma, 12-349) at a dilution of 1:4000. Membranes were washed three times in TBST, incubated with ECL detection reagent (Cytiva RPN2232) for 1 min, and imaged on a Bio-Rad ChemiDoc imager.

### Cloning, expression, and protein purification

Sequences of all proteins used in this study are in **Table S7**. To generate the GFP reporter plasmid, the native sequence of a Cerberus operon and the associated system’s promoter from *Vibrio parahaemolyticus* S082 were synthesized (GeneWiz ValueGene) and cloned via isothermal assembly into pLOCO3 upstream of a gene encoding GFP^68^.

For protein expression, *V. parahaemolyticus* CerA (IMG 2662261620) and CerB (IMG 2662261619) were amplified by PCR and cloned into UC Berkeley Macrolab vectors 2C-T (Addgene #29706; encoding an N-terminal TEV protease cleavable His_6_-MBP-tag), or 13S-A (Addgene #48323; no tag). Genes encoding *E. coli* ESBL60 CerC (NCBI WP_341816446.1) and CerB-GFP (NCBI WP_002431130.1) were codon-optimized for *E. coli* expression, synthesized (IDT), and cloned into UC Berkeley Macrolab vectors 2BC-T (Addgene #37236; C-terminal TEV protease cleavable His_6_-tag) and 2C-T, respectively. Point mutations were generated by PCR-based mutagenesis. For the isolated *E. coli* CerC peptidase domain, residues 1-183 were amplified by PCR and cloned into vector 2C-T. To generate the *E. coli* CerC Δloop constructs, residues 93-124 of WT CerC were replaced with a short linker (GSATSAG).

Proteins were expressed in *E. coli* T7 Express cells (New England Biolabs C2566). Cultures were grown in 1L of 2XYT medium at 37°C to A_600_ = 0.6, induced with 0.25 mM IPTG and moved to 20°C for 16-18 h. Cells were harvested by centrifugation and resuspended in buffer A (25 mM Tris, pH 8.5, 10% glycerol, 300 mM NaCl, 5 mM MgCl_2_, and 1 mM NaN_3_) plus 5 mM imidazole, 5 mM β-mercaptoethanol, and 5 µM ZnCl_2_ for purification of CerC. For high salt purification of *E. coli* CerC, buffer A contained 800 mM NaCl. Resuspensions were then lysed by sonication (Branson sonifier) and clarified by centrifugation, then passed over Ni-NTA agarose (Qiagen Ni-NTA Superflow) in resuspension buffer. The agarose was then washed twice with resuspension buffer, followed by wash buffer (buffer A plus 10 mM imidazole), and eluted in elution buffer (buffer A plus 400 mM imidazole). Proteins were then concentrated by ultrafiltration (Amicon Ultra; EMD Millipore). All proteins were passed over a Superdex 200 Increase size exclusion column (Cytiva) in SEC buffer (25 mM Tris, pH 8.5, 10% glycerol, 300 mM NaCl, 5 mM MgCl_2_, 1 mM DTT, and 1 mM NaN_3_). For cryo-EM samples, SEC buffer also contained 5% glycerol. Peak fractions were concentrated by ultrafiltration and stored at 4°C.

### Cryo-EM data collection and structure determination

For cryo-EM of apo CerC, purified protein was diluted to 10 µM in SEC buffer. 3.5 µL samples were applied to glow-discharged 300-mesh UltrAufoil grids (Electron Microscopy Sciences) in a Vitrobot Mark IV maintained at 4°C and 95% humidity (Thermo Fisher Scientific), blotted for 4 seconds with a blot force of 4.0, then vitrified in liquid ethane cooled by liquid nitrogen. The sample was imaged using a FEI Titan Krios G3 transmission electron microscope (Thermo Fisher Scientific) equipped with a K3 direct electron detector (Gatan) and a BioContinuum Energy Filter (Gatan) operating at 300 keV. The microscope was operated in EFTEM mode with a slit-width of 20 eV and using a 100 μm objective aperture. Automated data acquisition was performed using EPU (Thermo Fisher Scientific). Ten-second movies were collected at a magnification of 105,000x and a pixel size of 0.83 Å, with a total dose of 50 e^-^/Å^2^ distributed uniformly over 40 frames. In total, 1,572 movies were acquired with a realized defocus range of - 0.5 to -2.5 μm. After curation of micrographs based on ice thickness and contrast transfer function (CTF) fitting, 1,403 were accepted for particle picking.

For cryo-EM of DNA-bound CerC, purified protein was diluted to 10 µM in SEC buffer. 3.5 µL samples were applied to glow-discharged 300-mesh UltrAufoil grids (Electron Microscopy Sciences) in a Vitrobot Mark IV maintained at 4°C and 95% humidity (Thermo Fisher Scientific), blotted for 4 seconds with a blot force of 4.0, then vitrified in liquid ethane cooled by liquid nitrogen. The sample was imaged using a FEI Titan Krios G4 transmission electron microscope with Selectris X energy filter and Falcon 4 direct electron detector (Thermo Fisher Scientific) operating at 300 keV. The microscope was operated in EFTEM mode with a slit-width of 20 eV and using a 100 μm objective aperture. Automated data acquisition was performed using EPU (Thermo Fisher Scientific). Ten-second movies were collected at a magnification of 105,000x and a pixel size of 0.935 Å, with a total dose of 55 e^-^/Å^2^ distributed uniformly over 40 frames. In total, 3,636 movies were acquired with a realized defocus range of -0.5 to -2.5 μm. After curation of micrographs based on ice thickness and CTF fitting, 3,574 were accepted for particle picking.

All data processing was performed in cryoSPARC version 4 (Structura Biotechnology)^69^. For apo CerC, initial particle picking was done with the blob picker (637,034 particles), then particles were extracted and subjected to multiple rounds of 2D classification (259,701 particles). After *ab initio* reconstruction and heterogeneous refinement, a final set of 120,728 particles was subjected to non-uniform (NU) refinement, then reference- based motion correction and a final round of local refinement to yield a 3.07 Å resolution map (**Figure S7**). For DNA-bound CerC, initial blob-picked particles were sorted by 2D classification, *ab initio* reconstruction, and heterogeneous refinement, then 2D classes from the sorted particle set were used to generate templates for a second round of template-based particle picking (2,497,546 particles). After several rounds of 2D classification, *ab initio* reconstruction, and heterogeneous refinement, the particle stack was rebalanced to adjust for orientation bias, resulting in a final particle stack of 107,476 particles. This particle stack was subjected to NU refinement, then reference-based motion correction and a final round of local refinement to yield a 3.05 Å resolution map (**Figure S8**).

An initial model of CerC was generated using AlphaFold 3 ^70^. Models were docked into cryo-EM maps using ChimeraX^71^, followed by manual fitting and model rebuilding in Coot^72^ and real-space refinement in phenix.refine^73^ (**Table S3**).

### RNAP pull-down assay

To detect binding of *Vp* CerA to *E. coli* RNAP, 50 μL reactions of 10 μM purified MBP-CerA and/or 1 μM RNAP (purified as previously described)^74^ in 25 mM Tris-HCl pH 8.5, 200 mM NaCl, 5 mM MgCl_2_, 1 mM TCEP, and 0.01% Tween were assembled. 15 μL of the reaction was removed for the “input” gel sample followed by addition of 20 μL of amylose beads (New England Biolabs E8021S). Pull-down reactions were then incubated at 4°C for 1 hour before washing three times with reaction buffer and eluting with 5 μL of 0.1 M maltose. Elution was followed by addition of 30 μL of 2x SDS sample buffer and a western blot for the RNAP-ꞵ subunit using the protocol described above.

### DNA-binding assays

For characterization of DNA binding by fluorescence polarization (FP) assays, a 50-base pair double-stranded DNA was produced by annealing complementary oligos (sequence below), with the top strand 5ʹ 6-FAM labeled. FP reactions (30 μL) in buffer containing 25 mM Tris pH 8.5, 50 mM sodium glutamate, 5 mM MgCl_2_, 1 mM DTT, 5% glycerol, and 0.01% NP40 analog contained 50 nM DNA and the indicated MBP-CerA concentration. FP was read using a Tecan Spark plate reader, and binding data were analyzed with Graphpad Prism version 10 using a single-site binding model (Y = B_max_·X/ (*K_d_* + X) + Background). Top-strand sequence: 5ʹ-TAGTTGCGATTAAGTAAGTAAGACCAGAATGGTTCTGGCTGAAAACTAAA-3ʹ

### CerC helicase assays

For detecting CerC helicase activity *in vitro*, 12 µL reactions of 1 µM CerC, 2 mM ATP, and 200 ng of an 11 kb linearized plasmid were assembled in a reaction buffer containing 25 mM HEPES pH 7.5, 50 mM NaCl, 5 mM MgCl_2_, 5 µM ZnCl_2_, and 0.01% Tween-20. The reactions were incubated for 30 minutes at 37°C and quenched with 10 mM EDTA and 0.6% SDS. Following addition of 6x DNA loading dye, samples were loaded onto a 1% TAE-agarose gel, run for 30 minutes at 130 V, and imaged on a Bio-Rad ChemiDoc imager.

### CerC peptidase assays

To detect cleavage of CerB by CerC *in vitro*, 20 µL reactions of 1 µM CerC and 2 µM MBP-CerB-GFP in buffer 25 mM Tris pH 8.5, 50 mM NaCl, 5 mM MgCl_2_, and 5 µM ZnCl_2_ (cleavage buffer) were assembled with 2 mM ATP and/or 200 ng linearized plasmid DNA (described above) in the indicated samples. Assays performed with the isolated *E. coli* CerC peptidase domain contained 1 µM MBP-CerC-peptidase and 2 µM MBP-CerB-GFP in cleavage buffer with or without 1 µM of 22-nt polyT ssDNA. Reactions were incubated for 30 minutes at 37°C then added to 20 µL of 2x SDS sample buffer. 10 µL of each sample was loaded and separated by SDS-PAGE (TGX Stain-Free FastCast Acrylamide SDS-PAGE gel (Bio-Rad, 1610185)) and Coomassie-stained for visualization. For protein N-terminal sequencing by Edman degradation, peptidase reactions were separated by SDS-PAGE, transferred to a Bio-Rad Immun-Blot PVDF membrane, and stained with Coomassie Blue R-250. The C-terminal MBP-CerB-GFP cleavage product was excised, washed with methanol and deionized water, and loaded onto a Shimadzu PPSQ-53A instrument for sequence analysis (Iowa State University Protein Facility). The sample was subjected to five rounds of Edman degradation and the calculated amount yield (pmol) for each round was used to identify the likely sequence.

## ACKNOWLEDGEMENTS

The authors thank members of the Corbett lab for helpful discussions. The authors acknowledge funding from the National Institutes of Health (R35 GM122141 to K.D.C.; R35 GM142785 to T.V.M.; DP2 AT012346 to A.T.W.; T32 GM139795 to L.R.C., S.N.O., and B.Q.P.; F31 AI194603 to L.R.C.), the National Science Foundation (MCB 2516225 to T.V.M.), and the Burroughs Wellcome Fund (PATH Award 1186087 to A.T.W.). The authors acknowledge the use of the UC San Diego Cryo-EM facility, part of the Goeddel Family Technology Sandbox, along with the scientific and technical assistance of facility staff members Dr. Mariusz Matyszewski and Dr. Inga Kuschnerus.

## SUPPLEMENTAL INFORMATION

**Table S1.** Bub operons and associated transcriptional regulators. *See separate Excel file*

**Table S2.** Cerberus operons and associated immune/defense operons. *See separate Excel file*

**Table S3.** Cryoelectron microscopy data collection and refinement.

|  | **apo CerC** | **CerC-DNA** |
|---|---|---|
| **Data collection and processing** |  |  |
| Magnification | 105,000x | 105,000x |
| Voltage (kV) | 300 | 300 |
| Electron exposure (e<sup>-</sup>/Å<sup>2</sup>) | 50 | 55 |
| Defocus range (μm) | -0.5 to -2.5 | -0.5 to -2.5 |
| Pixel size (Å) | 0.83 | 0.935 |
| Symmetry imposed | N/A | N/A |
| Initial particle images (no.) | 637,034 | 2,497,546 |
| Final particle images (no.) | 120,728 | 107,476 |
| Map resolution (Å)<br>(FSC threshold) | 3.07 (0.143) | 3.05 (0.143) |
| **Refinement** |  |  |
| Model resolution (Å)<br>FSC threshold | 3.0 (0.143) | 3.0 (0.143) |
| Model composition |  |  |
| Non-hydrogen atoms | 8536 | 8865 |
| Protein residues | 1089 | 1072 |
| Nucleotide residues | 0 | 24 |
| Water | 0 | 0 |
| Ligands | 2 (Zn, SF4) | 2 (Zn, SF4) |
| *B* factors (Å<sup>2</sup>) |  |  |
| Protein | 103.1 | 88.2 |
| Nucleotide | N/A | 134.7 |
| Water | N/A | N/A |
| Ligand | 109.8 | 99.3 |
| R.m.s. deviations |  |  |
| Bond lengths (Å) | 0.003 | 0.004 |
| Bond angles (°) | 0.614 | 0.739 |
| **Validation** |  |  |
| MolProbity score | 1.42 | 1.51 |
| Clashscore | 4.45 | 5.77 |
| Poor rotamers (%) | 1.42 | 1.78 |
| Ramachandran plot |  |  |
| Favored (%) | 97.6 | 98.4 |
| Allowed (%) | 2.2 | 1.5 |
| Disallowed (%) | 0.2 | 0.1 |
| PDB ID | 38IU | 38JA |
| EMDB ID | EMD-78844 | EMD-78849 |

**Table S4.** Pairwise structure comparison of CerC helicase domain.

| Protein | Z-score | RMSD (Å) | Length of alignment | PDB:Chain |
|---|---|---|---|---|
| AdnB | 25.5 | 2.8 | 579 | 6PPJ:B |
| AdnA | 24.5 | 3.4 | 516 | 6PPJ:A |
| AddA | 24.5 | 3.5 | 626 | 3U44:A |
| AddB | 24.1 | 4.6 | 546 | 3U44:B |
| RecB | 20.1 | 4.0 | 564 | 3K70:B |
| RecC | 16.5 | 5.2 | 517 | 3K70:C |

**Table S5.** Pairwise structure comparison of CerC nuclease domain.

| Protein | Z-score | RMSD (Å) | Length of alignment | PDB:Chain |
|---|---|---|---|---|
| AdnA | 15.4 | 3.1 | 209 | 6PPJ:A |
| AdnB | 9.9 | 2.8 | 145 | 6PPJ:B |
| AddB | 9.4 | 3.7 | 222 | 3U44:B |
| RecB | 9.0 | 4.0 | 173 | 3K70:B |
| AddA | 8.0 | 3.8 | 151 | 3U44:A |
| RecC | 6.0 | 4.6 | 192 | 3K70:C |

**Table S6.** Orthrus operons. *See separate Excel file*

**Table S7.**
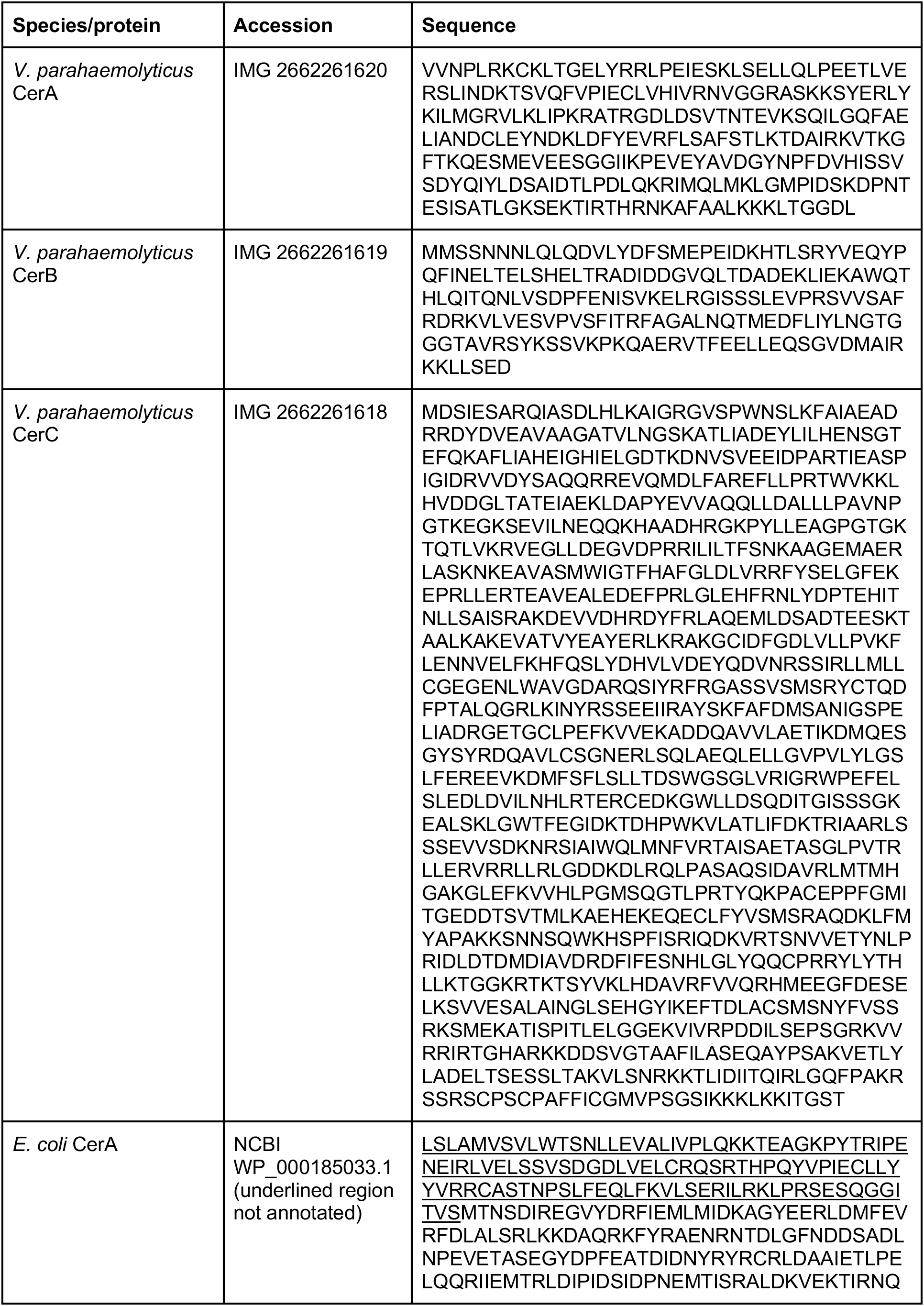
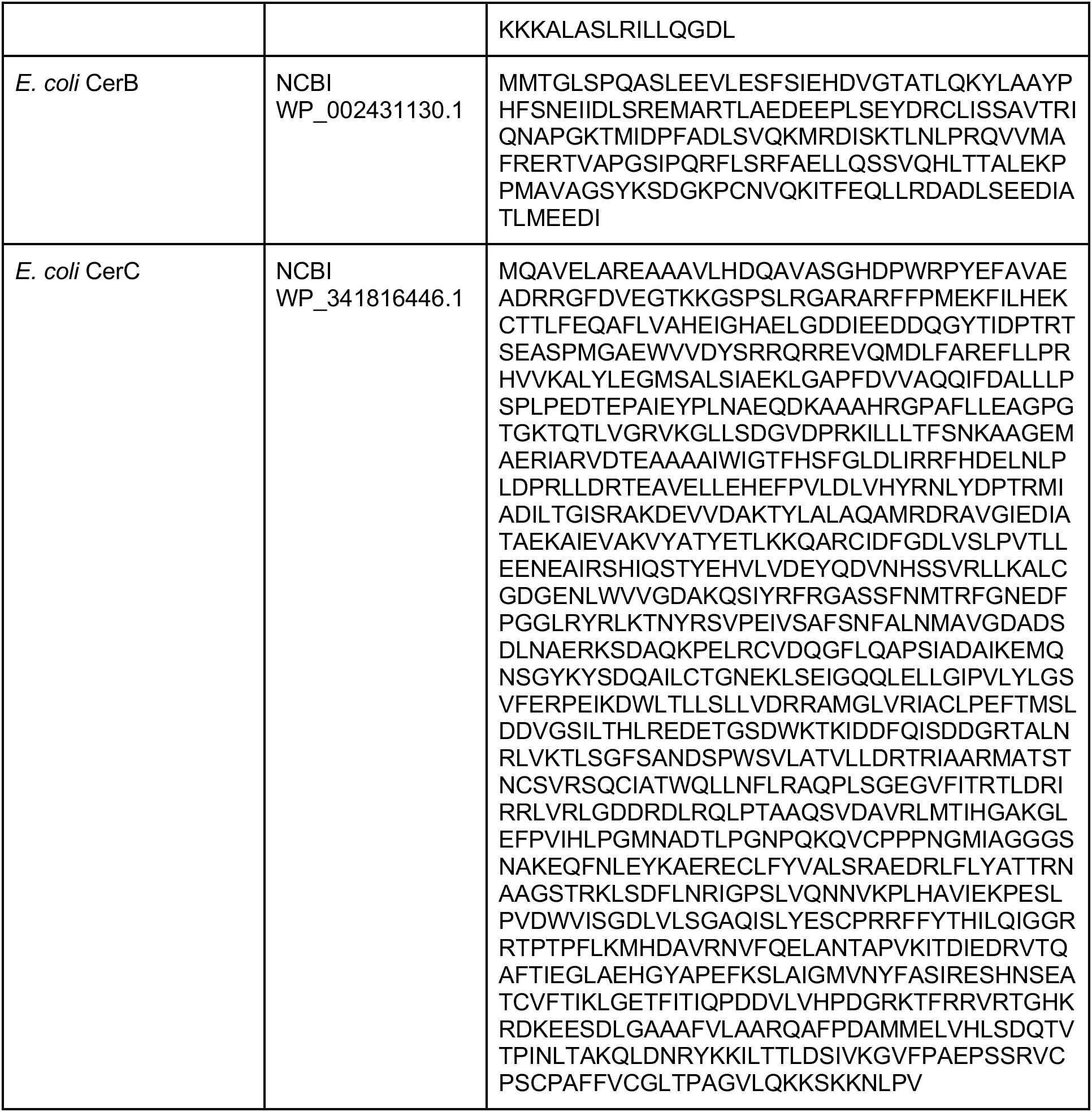
Proteins used in this study.

**Figure S1.**
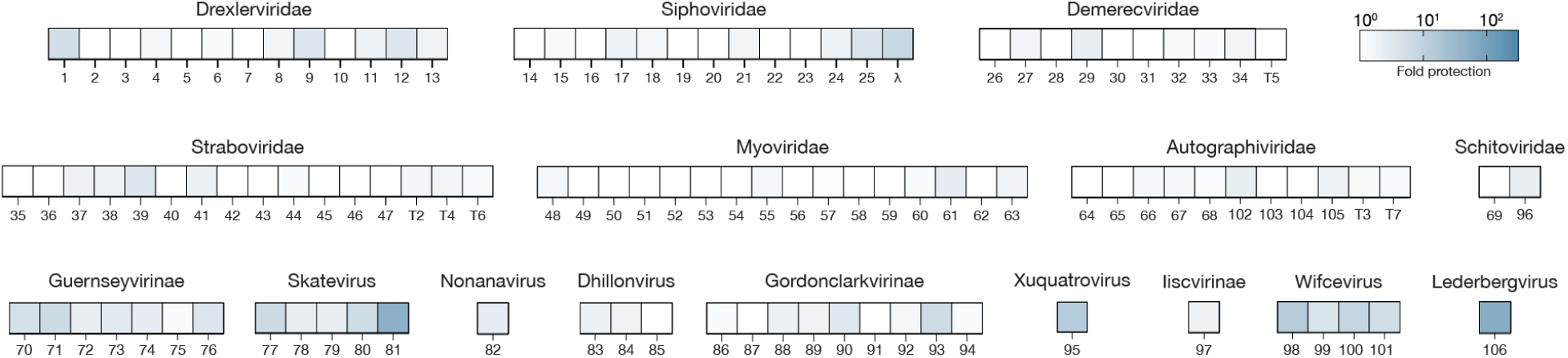
Plaque assays. Heat map of plaque assay results using the *V. parahaemolyticus* Cerberus-Bub locus in *E. coli*, from triplicate trials.

**Figure S2.**
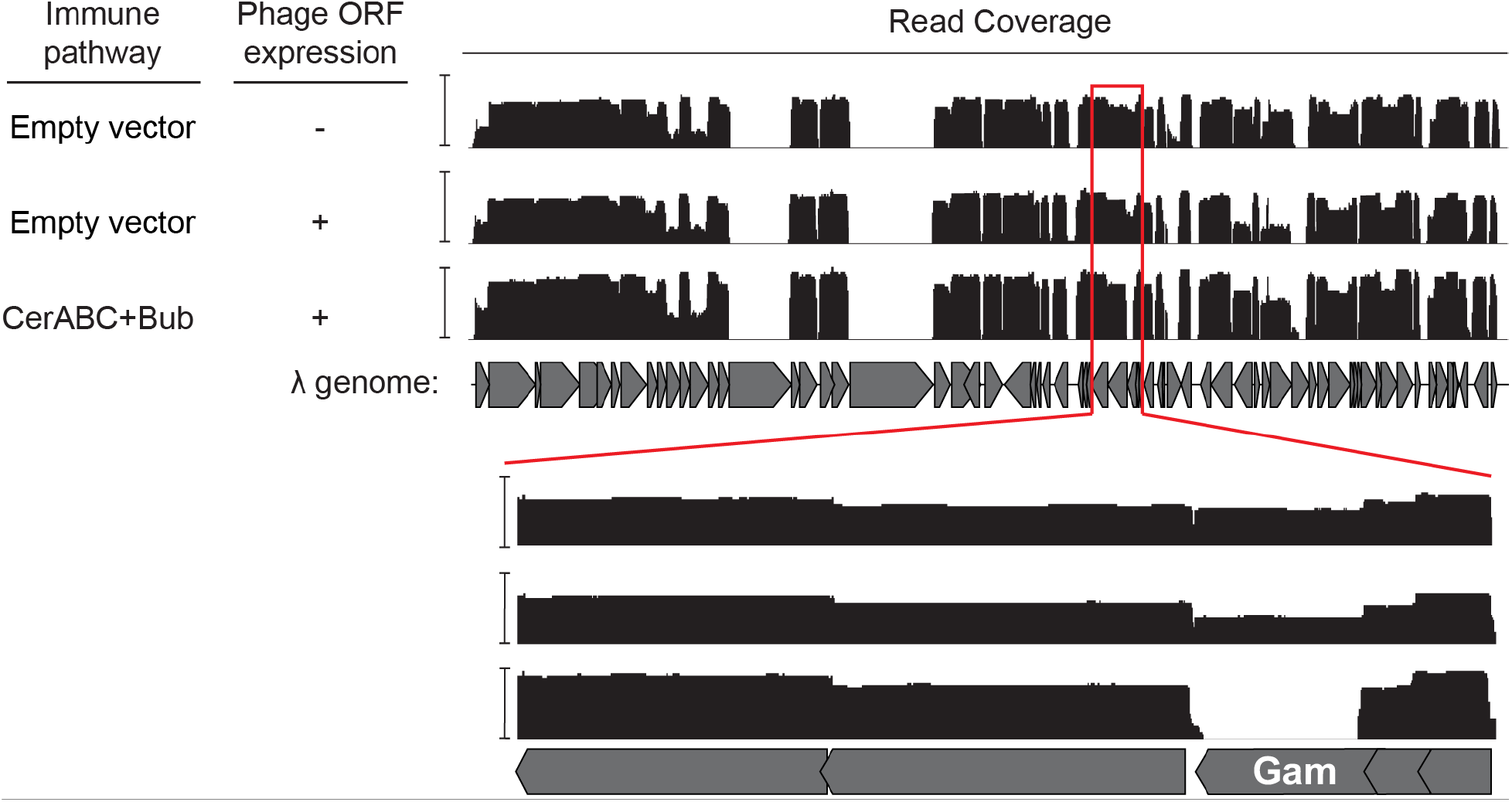
Phage ORF depletion assay. Read coverage from phage λ ORF depletion assay in the presence of the *V. parahaemolyticus* Cerberus-Bub locus. Y axes are shown as a log scale and are scaled to the maximum read coverage for each condition: 29,504 (Empty vector, no induction), 47,746 (Empty vector, induction), and 3,123 (Cerberus+Bub, induction).

**Figure S3.**
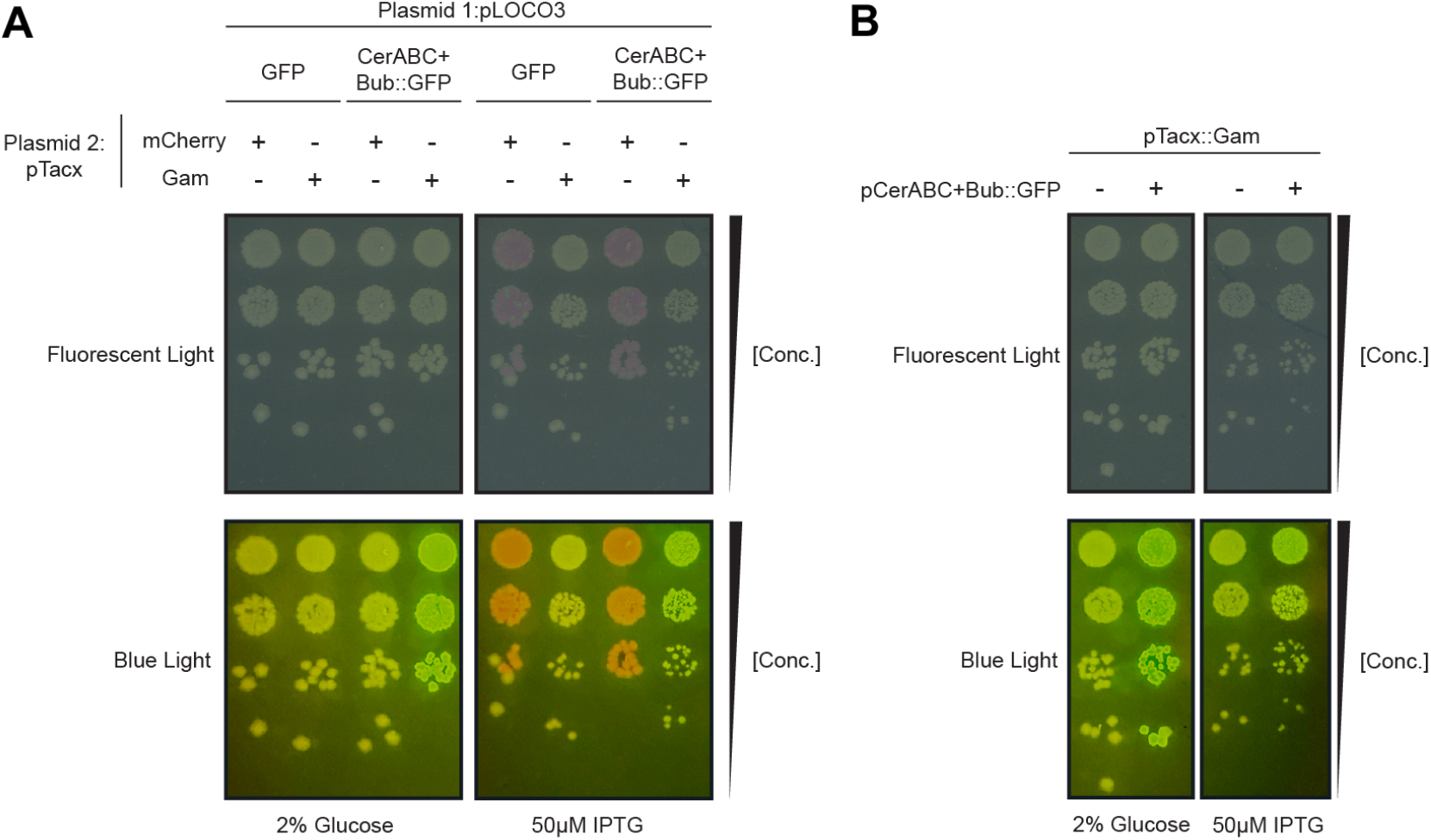
Expression of lambda Gam triggers Cerberus-mediated transcriptional activation. A. Bacterial dilutions imaged with white fluorescent light (top panels) or blue light to induce GFP/mCherry fluorescence (bottom), using a combination of plasmids as indicated. Plasmid 1 encodes either GFP alone or GFP under the control of Cerberus; plasmid 2 encodes either mCherry or λ Gam. B. Bacterial dilutions imaged as in panel (A), using a plasmid encoding GFP under the control of Cerberus and a second plasmid encoding λ Gam. 2% glucose: repression of λ Gam expression; 50 µM IPTG: induction of λ Gam expression.

**Figure S4.**
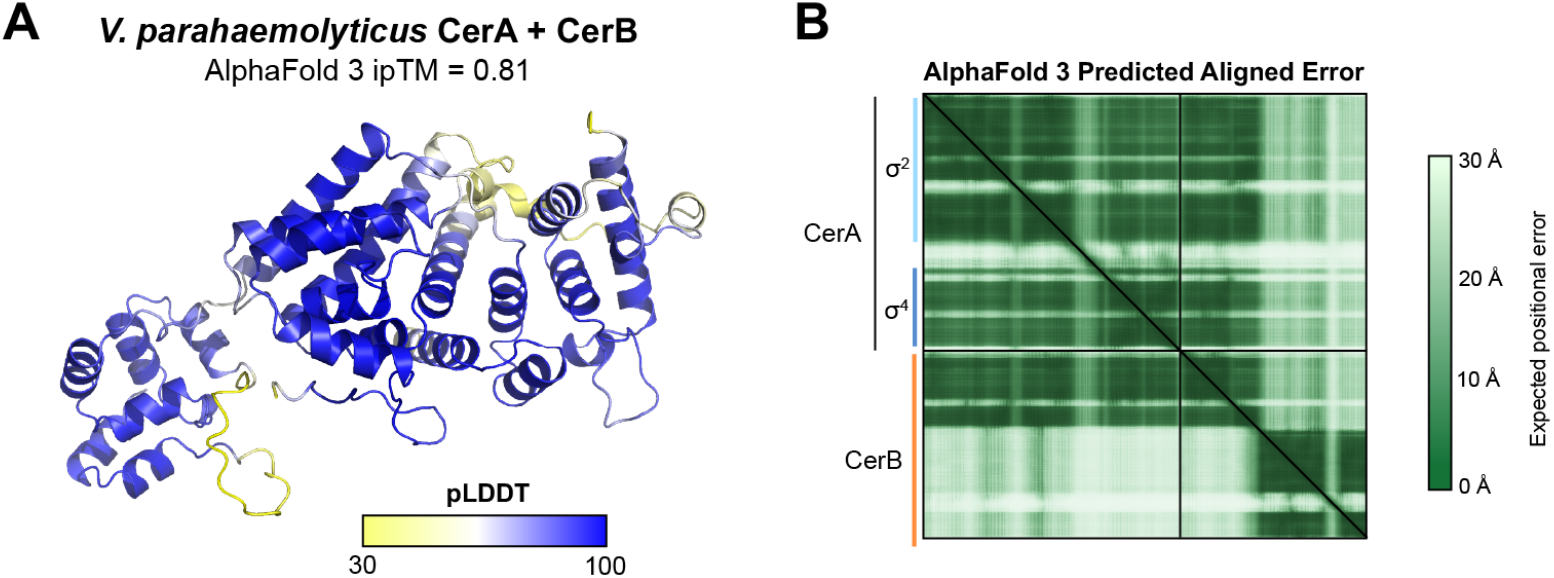
AlphaFold model of *V. parahaemolyticus* CerA-CerB. A. AlphaFold 3 model of a 1:1 complex of *V. parahaemolyticus* CerA and CerB (view as in Figure 2B), colored by local confidence (predicted local difference distance test; pLDDT). B. AlphaFold 3 predicted aligned error (PAE) plot for the prediction shown in panel (A).

**Figure S5.**
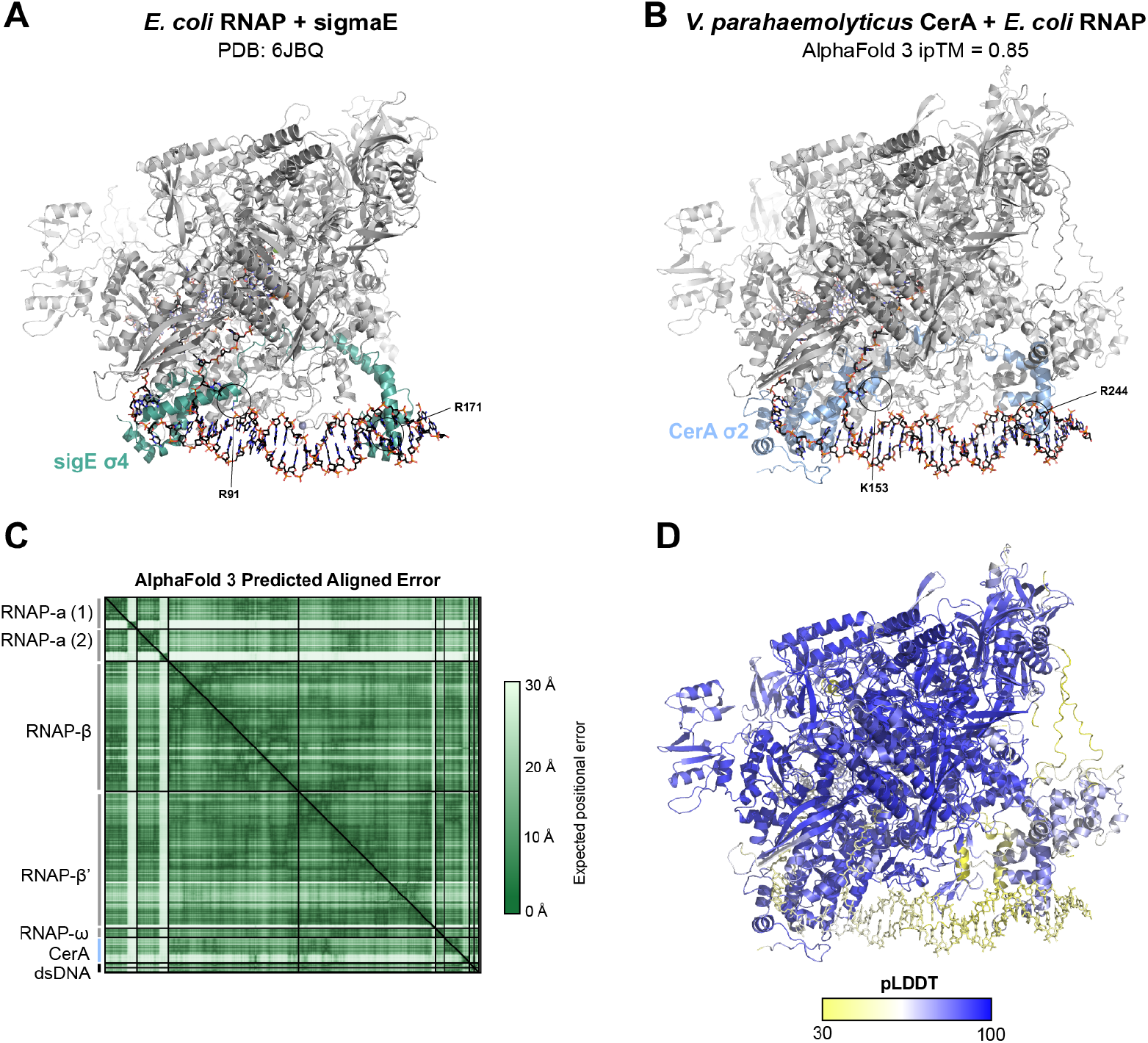
AlphaFold model of *V. parahaemolyticus* CerA bound to RNA polymerase and promoter DNA. A. Structure of *E. coli* RNA polymerase (gray) bound to the group IV sigma factor σ^E^ (green) and promoter DNA (PDB 6JBQ)^77^. Conserved DNA-interacting residues Arg91 and Arg171 are shown as sticks. B. AlphaFold 3 model of *E. coli* RNA polymerase (gray) bound to *V. parahaemolyticus* CerA (blue), with predicted DNA-interacting residues Lys153 and Arg244 shown as sticks. C. AlphaFold 3 predicted aligned error (PAE) plot for the prediction shown in panel (B). D. AlphaFold 3 model shown in panel (B), colored by local confidence (pLDDT).

**Figure S6.**
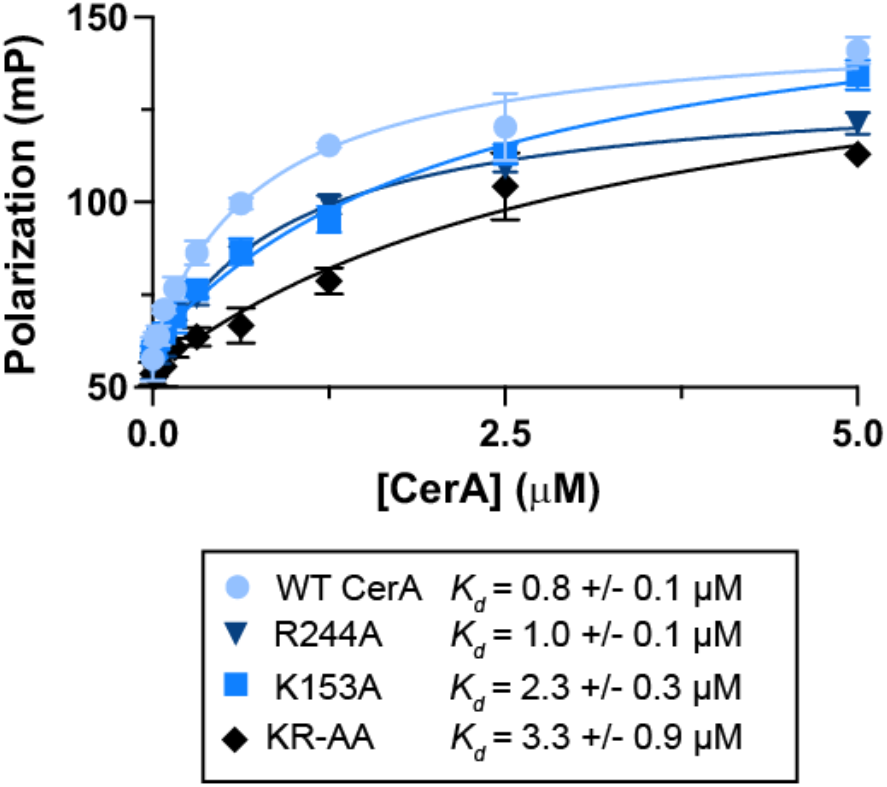
CerA-DNA binding. Fluorescence polarization DNA binding assay using the 50-bp double-stranded DNA probe shown in Figure 2D, and the indicated CerA proteins: WT (wild type), R224A, K153A, or K153A/R244A double mutant (KR-AA). *K_d_* values were calculated using a single-site binding model from triplicate data.

**Figure S7.**
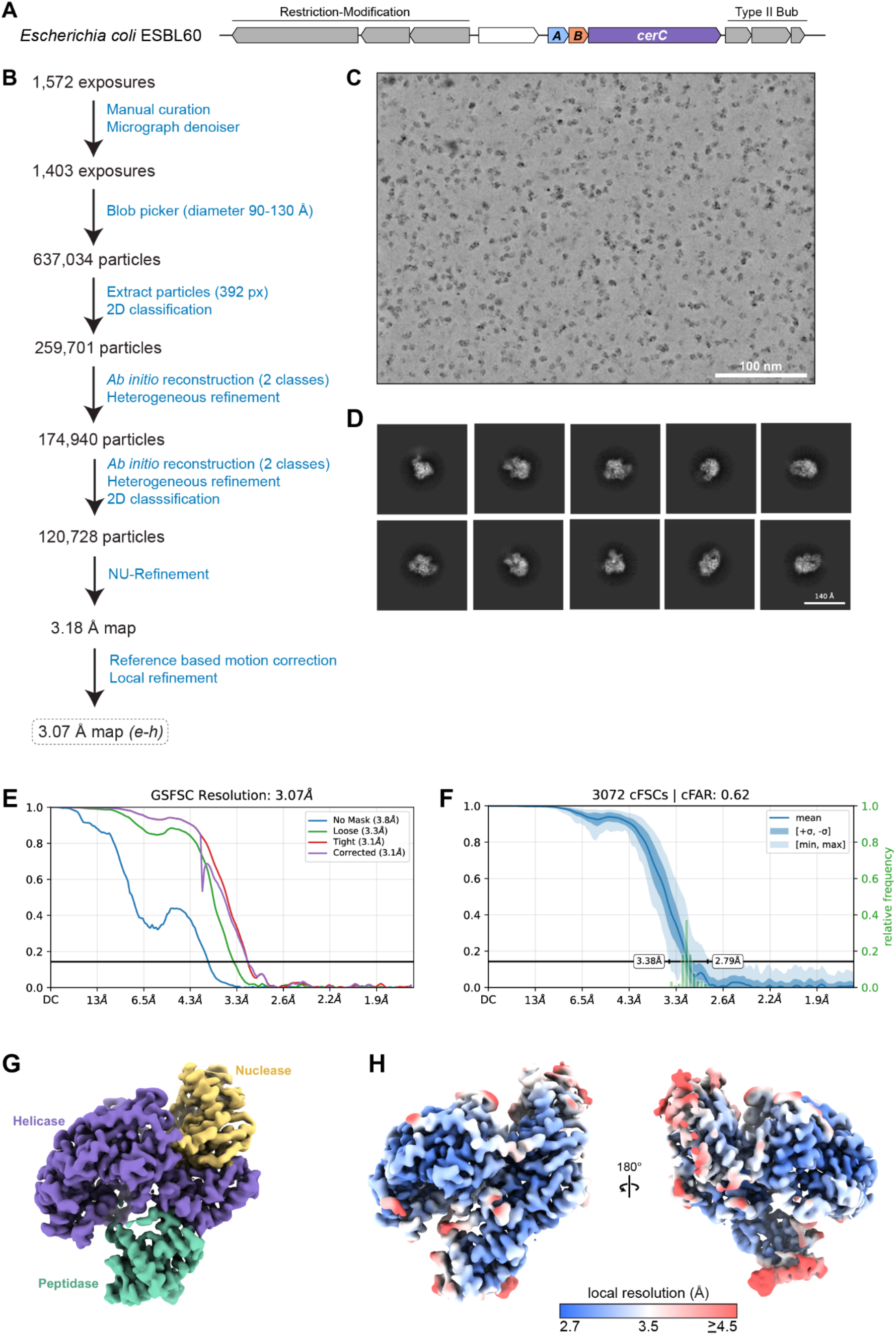
Cryo-EM workflow for apo CerC. A. Operon schematic of the *E. coli* ESBL60 Cerberus-Bub locus. B. Cryo-EM data collection and processing workflow for apo CerC. C. Denoised micrograph for apo CerC. D. Selected 2D classes for apo CerC. E. Fourier Shell Correlation (FSC) curve for the final refinement of apo CerC. F. Conical FSC Area Ratio (cFAR) curve for the final refinement of apo CerC. G. Refined cryo-EM map of apo CerC, with domains colored and labeled: peptidase green, helicase purple, and nuclease yellow. H. Two views of the refined cryo-EM map of apo CerC, colored by local resolution.

**Figure S8.**
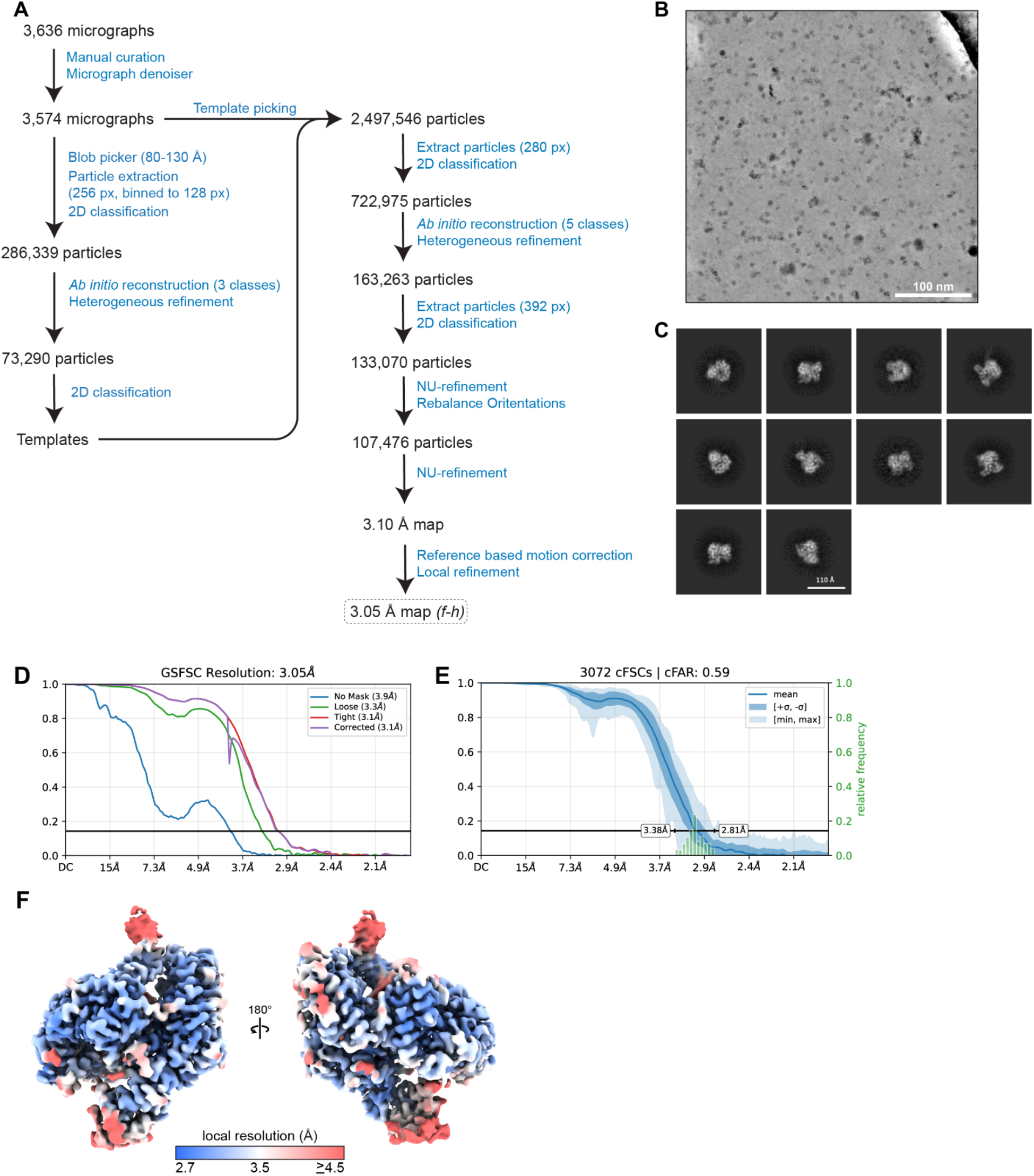
Cryo-EM workflow for DNA-bound CerC. A. Cryo-EM data collection and processing workflow for DNA-bound CerC. B. Denoised micrograph for DNA-bound CerC. C. Selected 2D classes for DNA-bound CerC. D. Fourier Shell Correlation (FSC) curve for the final refinement of DNA-bound CerC. E. Conical FSC Area Ratio (cFAR) curve for the final refinement of DNA-bound CerC. F. Two views of the refined cryo-EM map of DNA-bound CerC, colored by local resolution.

**Figure S9.**
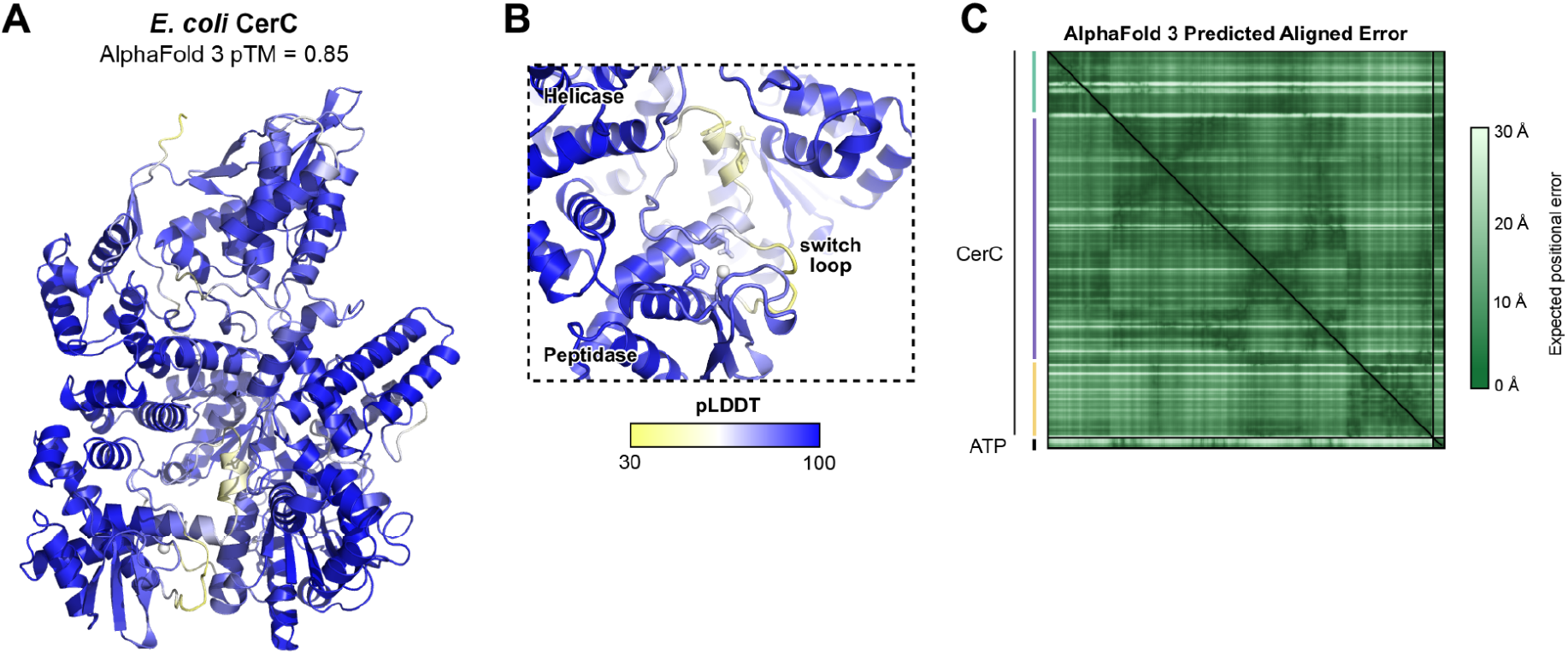
AlphaFold model of *E. coli* CerC. A. AlphaFold 3-predicted model of *E. coli* CerC, colored by local confidence (pLDDT). B. Closeup view of the peptidase active site with bound Zn^2+^ ion. C. AlphaFold 3 predicted aligned error (PAE) plot for the model shown in panel (A).

**Figure S10.**
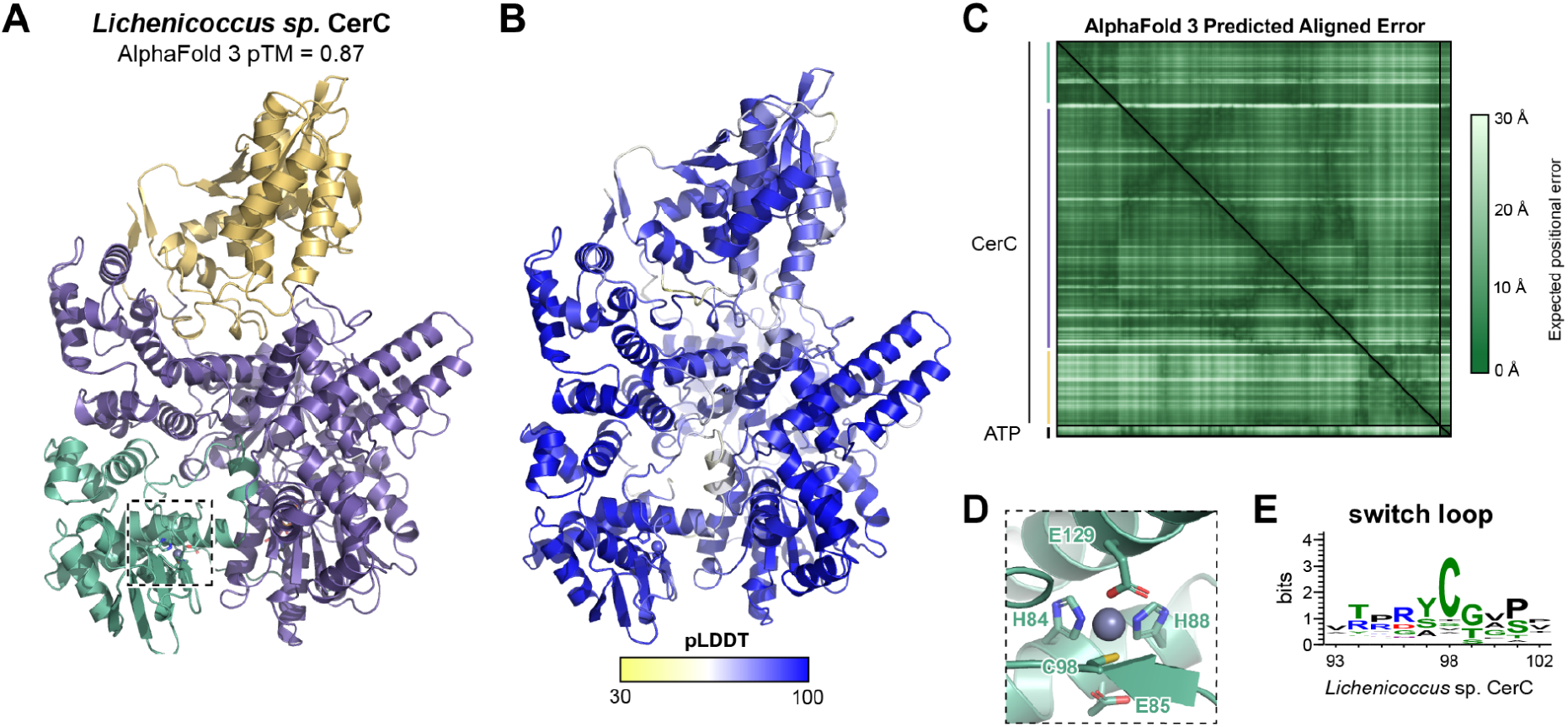
AlphaFold model of a CerC protein with a putative cysteine switch. A. AlphaFold 3 predicted model of *Lichenicoccus* sp. CerC (NCBI accession WP_428393404.1), colored by domain (peptidase green, helicase purple, nuclease yellow). B. The same model shown in panel (A), colored by local confidence (pLDDT). C. AlphaFold 3 predicted aligned error (PAE) plot for the model shown in panel (A). D. Closeup view of the peptidase active site with Zn^2+^ binding residues and the putative cysteine switch residue (Cys98) shown as sticks. E. Sequence logo for the switch loop region surrounding the putative cysteine switch. Cysteine is the most common residue at this position, found in 32% of sequences.

**Figure S11.**
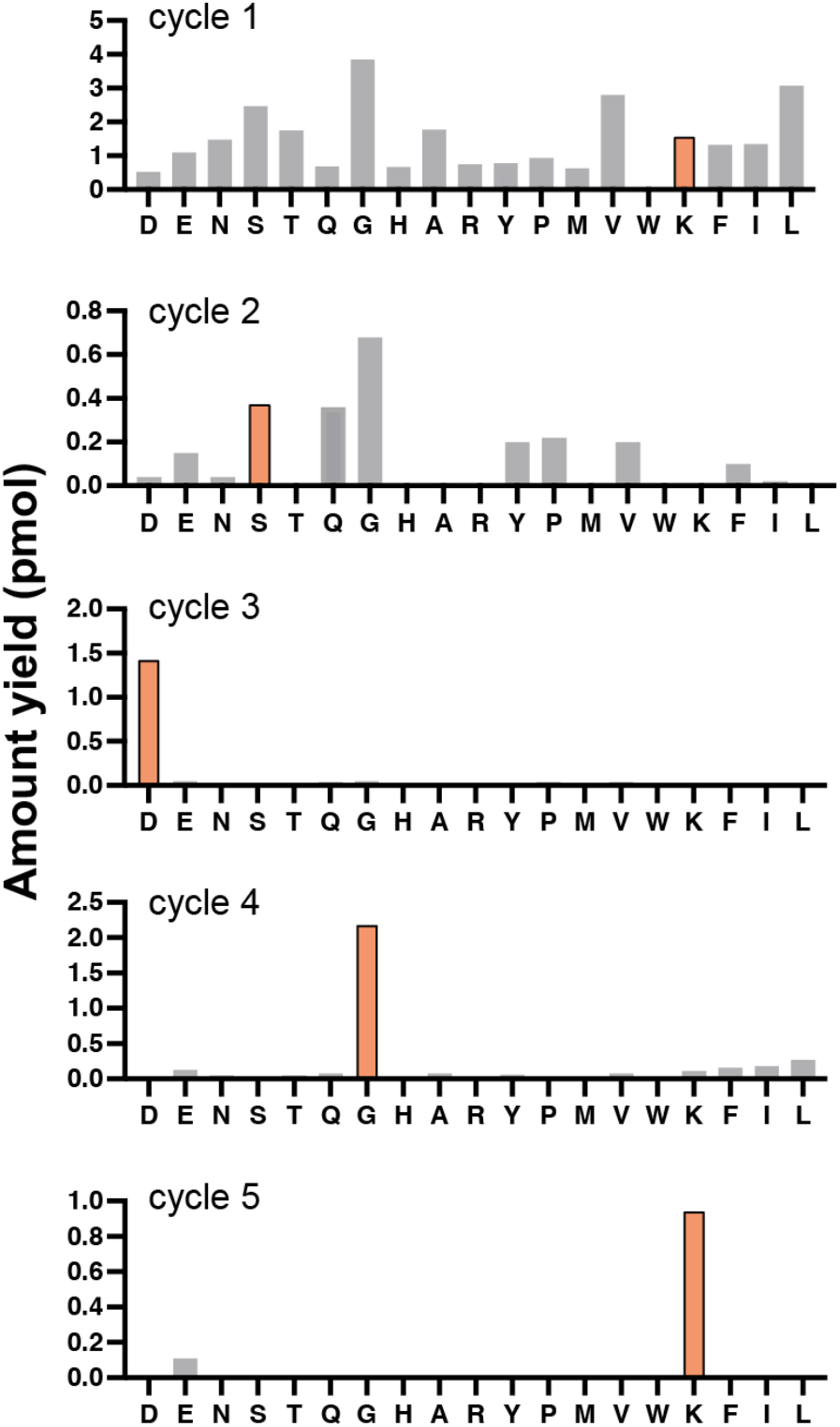
Edman degradation of CerB. Results of five cycles of Edman degradation of the C-terminal product of *E. coli* MBP-CerB-GFP cleavage from Figure 4D, graphed as amount yield (pmol). The correct amino acid from the inferred cleavage site on CerB (between Tyr157 and Lys158) is shown in orange: K (cycle 1), S (cycle 2), D (cycle 3), G (cycle 4), K (cycle 5).

**Figure S12.**
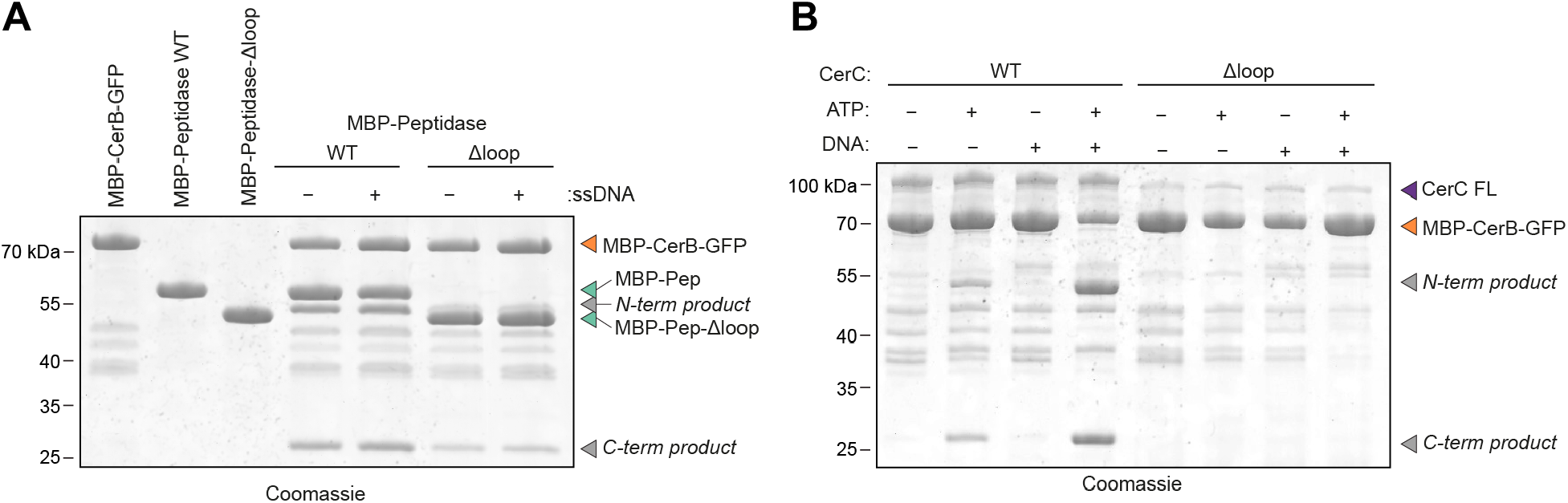
Effect of switch loop deletion on CerC peptidase activity. A. Coomassie-stained SDS-PAGE gel showing cleavage of MBP-CerB-GFP by the isolated CerC peptidase domain, wild type (WT) or Δloop mutant, in the presence and absence of single-stranded DNA (ssDNA). B. Coomassie-stained SDS-PAGE gel showing cleavage of MBP-CerB-GFP by full-length CerC, wild type (WT) or Δloop mutant, in the presence and absence of ATP and ssDNA.

**Figure S13.**
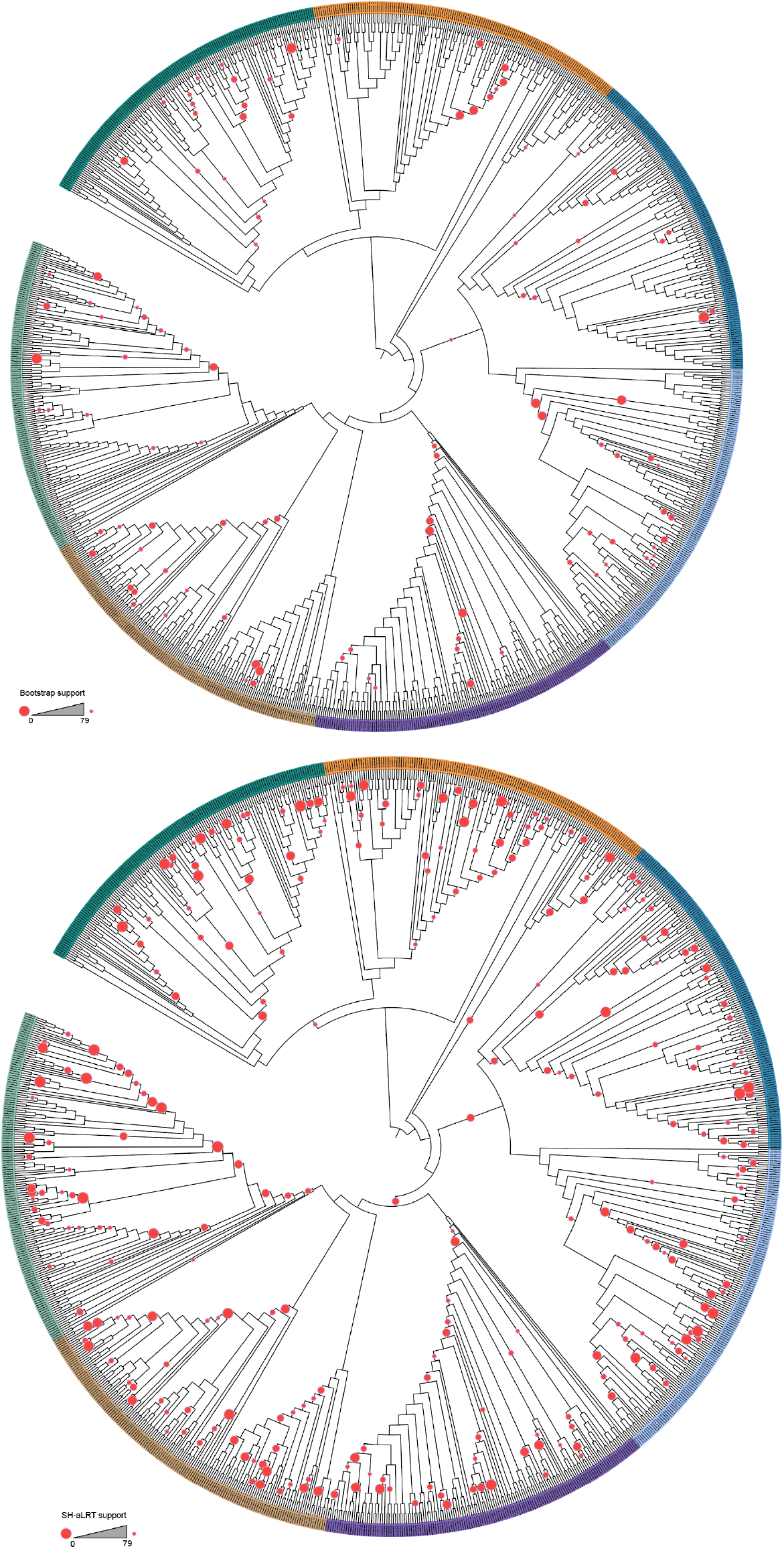
Support values for Helicase domain phylogenetic tree. A. Midpoint-rooted phylogenetic tree of the helicase domains of CerC and bacterial DNA repair proteins with red dots indicating Ultrafast bootstrap support values <80. B. Midpoint-rooted phylogenetic tree of the helicase domains of CerC and bacterial DNA repair proteins with red dots indicating SH-aLRT support values <80.

**Figure S14.**
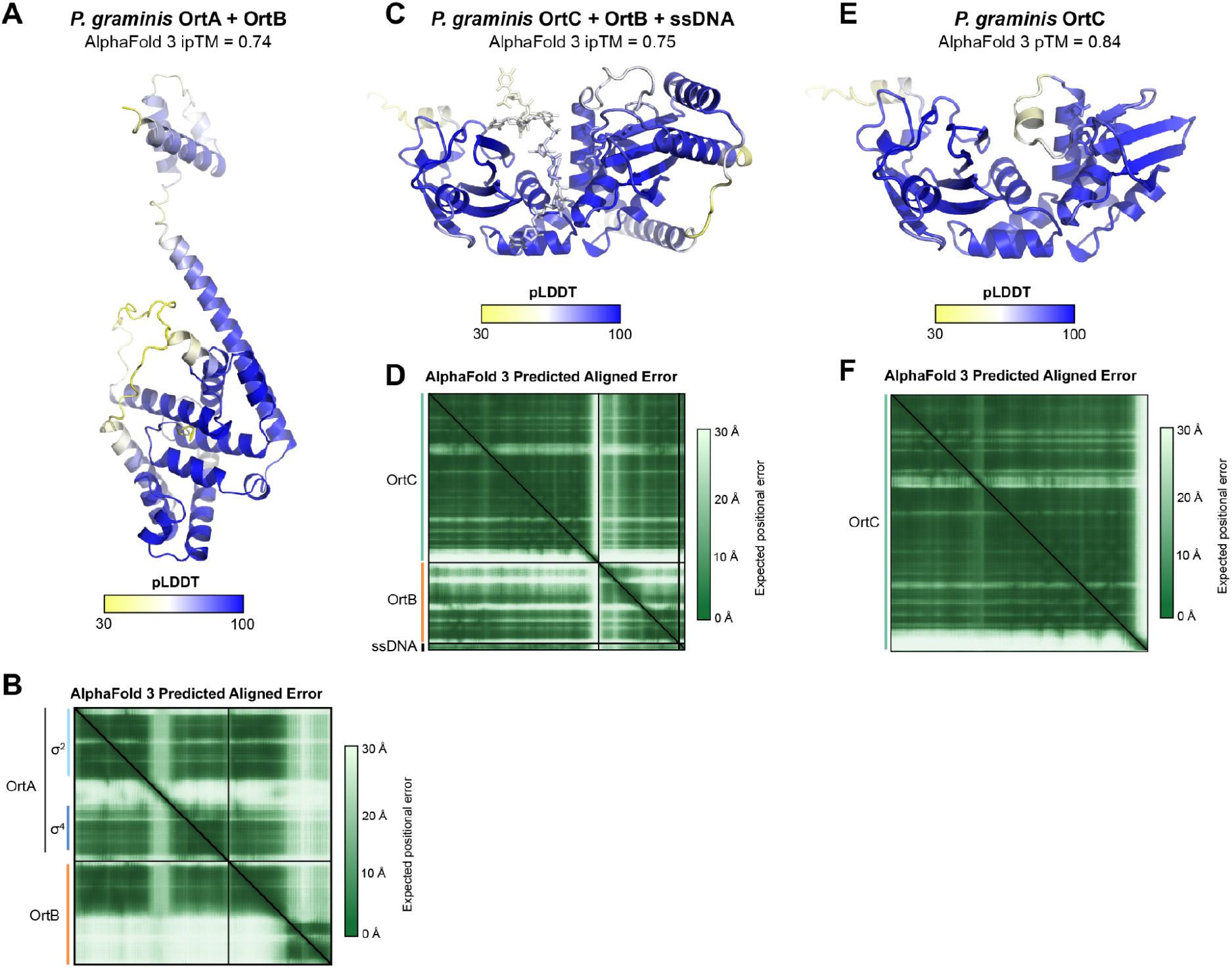
AlphaFold predictions of Orthrus proteins and complexes. A. AlphaFold 3 predicted structure of a *Paraburkholderia graminis* OrtA-OrtB complex, colored by local confidence (pLDDT). B. AlphaFold 3 predicted aligned error (PAE) for the prediction shown in panel (A). C. AlphaFold 3 predicted structure of a *P. graminis* OrtC-OrtB-ssDNA complex, colored by local confidence (pLDDT). D. AlphaFold 3 predicted aligned error (PAE) for the prediction shown in panel (C). E. AlphaFold 3 predicted structure of *P. graminis* OrtC, colored by local confidence (pLDDT). F. AlphaFold 3 predicted aligned error (PAE) for the prediction shown in panel (E).

## References

1. Bernheim, A., and Sorek, R. (2020). The pan-immune system of bacteria: antiviral defence as a community resource. Nat. Rev. Microbiol. 18, 113–119. 10.1038/s41579-019-0278-2.

2. Hampton, H.G., Watson, B.N.J., and Fineran, P.C. (2020). The arms race between bacteria and their phage foes. Nature 577, 327–336. 10.1038/s41586-019-1894-8.

3. Makarova, K.S., Wolf, Y.I., Snir, S., and Koonin, E.V. (2011). Defense Islands in Bacterial and Archaeal Genomes and Prediction of Novel Defense Systems. J. Bacteriol. 193, 6039–6056. 10.1128/jb.05535-11.

4. Juhas, M., van der Meer, J.R., Gaillard, M., Harding, R.M., Hood, D.W., and Crook, D.W. (2009). Genomic islands: tools of bacterial horizontal gene transfer and evolution. FEMS Microbiol. Rev. 33, 376–393. 10.1111/j.1574-6976.2008.00136.x.

5. Doron, S., Melamed, S., Ofir, G., Leavitt, A., Lopatina, A., Keren, M., Amitai, G., and Sorek, R. (2018). Systematic discovery of antiphage defense systems in the microbial pangenome. Science 359, eaar4120. 10.1126/science.aar4120.

6. Gao, Y., Cao, D., Zhu, J., Feng, H., Luo, X., Liu, S., Yan, X.-X., Zhang, X., and Gao, P. (2020). Structural insights into assembly, operation and inhibition of a type I restriction–modification system. Nat. Microbiol. 5, 1107–1118. 10.1038/s41564-020-0731-z.

7. Johnson, A.G., Wein, T., Mayer, M.L., Duncan-Lowey, B., Yirmiya, E., Oppenheimer-Shaanan, Y., Amitai, G., Sorek, R., and Kranzusch, P.J. (2022). Bacterial gasdermins reveal an ancient mechanism of cell death. Science 375, 221–225. 10.1126/science.abj8432.

8. Millman, A., Melamed, S., Leavitt, A., Doron, S., Bernheim, A., Hör, J., Garb, J., Bechon, N., Brandis, A., Lopatina, A., et al. (2022). An expanded arsenal of immune systems that protect bacteria from phages. Cell Host Microbe 30, 1556–1569.e5. 10.1016/j.chom.2022.09.017.

9. Lopatina, A., Tal, N., and Sorek, R. (2020). Abortive Infection: Bacterial Suicide as an Antiviral Immune Strategy. Annu. Rev. Virol. 7, 371–384. 10.1146/annurev-virology-011620-040628.

10. Tamulaitiene, G., Sabonis, D., Sasnauskas, G., Ruksenaite, A., Silanskas, A., Avraham, C., Ofir, G., Sorek, R., Zaremba, M., and Siksnys, V. (2024). Activation of Thoeris antiviral system via SIR2 effector filament assembly. Nature 627, 431–436. 10.1038/s41586-024-07092-x.

11. Saxton, D.S., and Laub, M.T. (2026). How bacterial immune systems sense phage infection. Nat. Rev. Microbiol., 1–13. 10.1038/s41579-026-01376-x.

12. Høyland-Kroghsbo, N.M., Paczkowski, J., Mukherjee, S., Broniewski, J., Westra, E., Bondy-Denomy, J., and Bassler, B.L. (2017). Quorum sensing controls the Pseudomonas aeruginosa CRISPR-Cas adaptive immune system. Proc. Natl. Acad. Sci. U. S. A. 114, 131–135. 10.1073/pnas.1617415113.

13. Patterson, A.G., Jackson, S.A., Taylor, C., Evans, G.B., Salmond, G.P.C., Przybilski, R., Staals, R.H.J., and Fineran, P.C. (2016). Quorum Sensing Controls Adaptive Immunity through the Regulation of Multiple CRISPR-Cas Systems. Mol. Cell 64, 1102–1108. 10.1016/j.molcel.2016.11.012.

14. Negri, A., Werbowy, O., Wons, E., Dersch, S., Hinrichs, R., Graumann, P.L., and Mruk, I. (2021). Regulator-dependent temporal dynamics of a restriction-modification system’s gene expression upon entering new host cells: single-cell and population studies. Nucleic Acids Res. 49, 3826–3840. 10.1093/nar/gkab183.

15. Lau, R.K., Enustun, E., Gu, Y., Nguyen, J.V., and Corbett, K.D. (2022). A conserved signaling pathway activates bacterial CBASS immune signaling in response to DNA damage. EMBO J. 41, e111540. 10.15252/embj.2022111540.

16. Luyten, Y.A., Hausman, D.E., Young, J.C., Doyle, L.A., Higashi, K.M., Ubilla-Rodriguez, N.C., Lambert, A.R., Arroyo, C.S., Forsberg, K.J., Morgan, R.D., et al. (2022). Identification and characterization of the WYL BrxR protein and its gene as separable regulatory elements of a BREX phage restriction system. Nucleic Acids Res. 50, 5171–5190. 10.1093/nar/gkac311.

17. Blankenchip, C.L., Nguyen, J.V., Lau, R.K., Ye, Q., Gu, Y., and Corbett, K.D. (2022). Control of bacterial immune signaling by a WYL domain transcription factor. Nucleic Acids Res. 50, 5239–5250. 10.1093/nar/gkac343.

18. Blankenchip, C.L., and Corbett, K.D. (2024). Bacterial WYL domain transcriptional repressors sense single-stranded DNA to control gene expression. Nucleic Acids Res., gkae1101. 10.1093/nar/gkae1101.

19. Chambers, L.R., Rani, P., Min, R.K., Villa, E., and Corbett, K.D. (2026). A DNA damage-activated kinase phosphorylates a transcriptional repressor to control bacterial immune pathway expression. EMBO J. 45, 5079–5100. 10.1038/s44318-026-00831-y.

20. Picton, D.M., Harling-Lee, J.D., Duffner, S.J., Went, S.C., Morgan, R.D., Hinton, J.C.D., and Blower, T.R. (2022). A widespread family of WYL-domain transcriptional regulators co-localizes with diverse phage defence systems and islands. Nucleic Acids Res. 50, 5191–5207. 10.1093/nar/gkac334.

21. Gong, M., Ye, Q., Gu, Y., Chambers, L.R., Bobkov, A.A., Arakawa, N.K., Matyszewski, M., and Corbett, K.D. (2025). Structural diversity and oligomerization of bacterial ubiquitin-like proteins. Structure. 33, 1016–1026.e4. 10.1016/j.str.2025.03.011.

22. Ye, Q., Gong, M., Cai, J., Chambers, L.R., Zhou, H., Suhandynata, R.T., and Corbett, K.D. (2026). Mechanistic basis for protein conjugation in a diverged bacterial ubiquitination pathway. Nat. Struct. Mol. Biol. 33, 71–83. 10.1038/s41594-025-01696-1.

23. Ledvina, H.E., Ye, Q., Gu, Y., Sullivan, A.E., Quan, Y., Lau, R.K., Zhou, H., Corbett, K.D., and Whiteley, A.T. (2023). An E1–E2 fusion protein primes antiviral immune signalling in bacteria. Nature 616, 319–325. 10.1038/s41586-022-05647-4.

24. Cohen, D., Melamed, S., Millman, A., Shulman, G., Oppenheimer-Shaanan, Y., Kacen, A., Doron, S., Amitai, G., and Sorek, R. (2019). Cyclic GMP-AMP signalling protects bacteria against viral infection. Nature 574, 691–695. 10.1038/s41586-019-1605-5.

25. Chambers, L.R., Ye, Q., Cai, J., Gong, M., Ledvina, H.E., Zhou, H., Whiteley, A.T., Suhandynata, R.T., and Corbett, K.D. (2024). A eukaryotic-like ubiquitination system in bacterial antiviral defence. Nature, 1–7. 10.1038/s41586-024-07730-4.

26. Hör, J., Wolf, S.G., and Sorek, R. (2024). Bacteria conjugate ubiquitin-like proteins to interfere with phage assembly. Nature, 1–7. 10.1038/s41586-024-07616-5.

27. Connelly, J.C., Kirkham, L.A., and Leach, D.R. (1998). The SbcCD nuclease of Escherichia coli is a structural maintenance of chromosomes (SMC) family protein that cleaves hairpin DNA. Proc. Natl. Acad. Sci. U. S. A. 95, 7969–7974. 10.1073/pnas.95.14.7969.

28. Humolli, D., Piel, D., Maffei, E., Heyer, Y., Agustoni, E., Shaidullina, A., Willi, L., Imwinkelried, P., Estermann, F., Cuénod, A., et al. (2025). Completing the BASEL phage collection to unlock hidden diversity for systematic exploration of phage– host interactions. PLOS Biol. 23, e3003063. 10.1371/journal.pbio.3003063.

29. Nagy, T.A., Gersabeck, G.W., Conte, A.N., and Whiteley, A.T. (2026). A phage protein screen identifies triggers of the bacterial innate immune system. Nat. Microbiol. 11, 597–609. 10.1038/s41564-025-02239-6.

30. Baba, T., Ara, T., Hasegawa, M., Takai, Y., Okumura, Y., Baba, M., Datsenko, K.A., Tomita, M., Wanner, B.L., and Mori, H. (2006). Construction of Escherichia coli K-12 in-frame, single-gene knockout mutants: the Keio collection. Mol. Syst. Biol. 2, 2006.0008. 10.1038/msb4100050.

31. Campbell, E.A., Tupy, J.L., Gruber, T.M., Wang, S., Sharp, M.M., Gross, C.A., and Darst, S.A. (2003). Crystal Structure of Escherichia coli σE with the Cytoplasmic Domain of Its Anti-σ RseA. Mol. Cell 11, 1067–1078. 10.1016/S1097-2765(03)00148-5.

32. Treviño-Quintanilla, L.G., Freyre-González, J.A., and Martínez-Flores, I. (2013). Anti-Sigma Factors in E. coli: Common Regulatory Mechanisms Controlling Sigma Factors Availability. Curr. Genomics 14, 378–387. 10.2174/1389202911314060007.

33. Ades, S.E. (2004). Control of the alternative sigma factor σE in *Escherichia coli*. Curr. Opin. Microbiol. 7, 157–162. 10.1016/j.mib.2004.02.010.

34. Vujičić-Žagar, A., Dulermo, R., Le Gorrec, M., Vannier, F., Servant, P., Sommer, S., De Groot, A., and Serre, L. (2009). Crystal Structure of the IrrE Protein, a Central Regulator of DNA Damage Repair in Deinococcaceae. J. Mol. Biol. 386, 704–716. 10.1016/j.jmb.2008.12.062.

35. Dillingham, M.S., and Kowalczykowski, S.C. (2008). RecBCD Enzyme and the Repair of Double-Stranded DNA Breaks. Microbiol. Mol. Biol. Rev. MMBR 72, 642– 671. 10.1128/MMBR.00020-08.

36. Krajewski, W.W., Fu, X., Wilkinson, M., Cronin, N.B., Dillingham, M.S., and Wigley, D.B. (2014). Structural basis for translocation by AddAB helicase–nuclease and its arrest at χ sites. Nature 508, 416–419. 10.1038/nature13037.

37. Sinha, K.M., Unciuleac, M.-C., Glickman, M.S., and Shuman, S. (2009). AdnAB: a new DSB-resecting motor–nuclease from mycobacteria. Genes Dev. 23, 1423– 1437. 10.1101/gad.1805709.

38. Jia, N., Unciuleac, M.C., Xue, C., Greene, E.C., Patel, D.J., and Shuman, S. (2019). Structures and single-molecule analysis of bacterial motor nuclease AdnAB illuminate the mechanism of DNA double-strand break resection. Proc. Natl. Acad. Sci. 116, 24507–24516. 10.1073/pnas.1913546116.

39. Warren, G.M., Jason de la Cruz, M., Goldgur, Y., and Shuman, S. (2026). Structure of an initiation complex of mycobacterial helicase-nuclease AdnAB at a blunt double-strand break reveals local melting that engages the 3′ tracking strand at the ratchet pawl of the helicase motor. Nucleic Acids Res. 54, gkag243. 10.1093/nar/gkag243.

40. Unciuleac, M.-C., Meir, A., Xue, C., Warren, G.M., Greene, E.C., and Shuman, S. (2021). Clutch mechanism of chemomechanical coupling in a DNA resecting motor nuclease. Proc. Natl. Acad. Sci. 118, e2023955118. 10.1073/pnas.2023955118.

41. Lu, H., Chen, Z., Xie, T., Zhong, S., Suo, S., Song, S., Wang, L., Xu, H., Tian, B., Zhao, Y., et al. (2024). The Deinococcus protease PprI senses DNA damage by directly interacting with single-stranded DNA. Nat. Commun. 15, 1892. 10.1038/s41467-024-46208-9.

42. Wigley, D.B. (2013). Bacterial DNA repair: recent insights into the mechanism of RecBCD, AddAB and AdnAB. Nat. Rev. Microbiol. 11, 9–13. 10.1038/nrmicro2917.

43. Holm, L., Laiho, A., Törönen, P., and Salgado, M. (2023). DALI shines a light on remote homologs: One hundred discoveries. Protein Sci. 32, e4519. 10.1002/pro.4519.

44. Bose, B., and Grossman, A.D. (2011). Regulation of Horizontal Gene Transfer in *Bacillus subtilis* by Activation of a Conserved Site-Specific Protease. J. Bacteriol. 193, 22–29. 10.1128/JB.01143-10.

45. Bose, B., Auchtung, J.M., Lee, C.A., and Grossman, A.D. (2008). A conserved anti-repressor controls horizontal gene transfer by proteolysis. Mol. Microbiol. 70, 570–582. 10.1111/j.1365-2958.2008.06414.x.

46. Ludanyi, M., Blanchard, L., Dulermo, R., Brandelet, G., Bellanger, L., Pignol, D., Lemaire, D., and De Groot, A. (2014). Radiation response in *Deinococcus deserti*: IrrE is a metalloprotease that cleaves repressor protein DdrO. Mol. Microbiol. 94, 434–449. 10.1111/mmi.12774.

47. Singh, M., Singh, C., Nair, A.V., Ahmad, I., Sharma, A., Bhasin, M., Jain, V., Singh, R., and Thakur, K.G. (2026). Structural and functional characterization of VapBC52 toxin–antitoxin system from *Mycobacterium tuberculosis*. Nucleic Acids Res. 54, gkag611. 10.1093/nar/gkag611.

48. Hayes, F., and Kędzierska, B. (2014). Regulating Toxin-Antitoxin Expression: Controlled Detonation of Intracellular Molecular Timebombs. Toxins 6, 337–358. 10.3390/toxins6010337.

49. Jurėnas, D., Fraikin, N., Goormaghtigh, F., and Van Melderen, L. (2022). Biology and evolution of bacterial toxin–antitoxin systems. Nat. Rev. Microbiol. 20, 335–350. 10.1038/s41579-021-00661-1.

50. Payne, L.J., Todeschini, T.C., Wu, Y., Perry, B.J., Ronson, C.W., Fineran, P.C., Nobrega, F.L., and Jackson, S.A. (2021). Identification and classification of antiviral defence systems in bacteria and archaea with PADLOC reveals new system types. Nucleic Acids Res. 49, 10868–10878. 10.1093/nar/gkab883.

51. Katoh, K., Rozewicki, J., and Yamada, K.D. (2019). MAFFT online service: multiple sequence alignment, interactive sequence choice and visualization. Brief. Bioinform. 20, 1160–1166. 10.1093/bib/bbx108.

52. Waterhouse, A.M., Procter, J.B., Martin, D.M.A., Clamp, M., and Barton, G.J. (2009). Jalview Version 2—a multiple sequence alignment editor and analysis workbench. Bioinformatics 25, 1189–1191. 10.1093/bioinformatics/btp033.

53. Crooks, G.E., Hon, G., Chandonia, J.M., and Brenner, S.E. (2004). WebLogo: a sequence logo generator. Genome Res. 14, 1188–1190. 10.1101/gr.849004.

54. Mutte, S.K., Barendse, P., Ugarte, P.B., and Swarts, D.C. (2025). Distribution of bacterial DNA repair proteins and their co-occurrence with immune systems. Cell Rep. 44, 115110. 10.1016/j.celrep.2024.115110.

55. Altschul, S.F., Gish, W., Miller, W., Myers, E.W., and Lipman, D.J. (1990). Basic local alignment search tool. J. Mol. Biol. 215, 403–410. 10.1016/S0022-2836(05)80360-2.

56. Shen, W., Le, S., Li, Y., and Hu, F. (2016). SeqKit: A Cross-Platform and Ultrafast Toolkit for FASTA/Q File Manipulation. PLOS ONE 11, e0163962. 10.1371/journal.pone.0163962.

57. Rajković, A., Beracochea, M., Rogers, A.B., Eddy, S.R., Carter, N.P., and Finn, R.D. (2026). HMMER web server: 2026 update. Nucleic Acids Res. 54, W272– W278. 10.1093/nar/gkag373.

58. Katoh, K., Misawa, K., Kuma, K., and Miyata, T. (2002). MAFFT: a novel method for rapid multiple sequence alignment based on fast Fourier transform. Nucleic Acids Res. 30, 3059–3066. 10.1093/nar/gkf436.

59. Katoh, K., and Standley, D.M. (2013). MAFFT Multiple Sequence Alignment Software Version 7: Improvements in Performance and Usability. Mol. Biol. Evol. 30, 772–780. 10.1093/molbev/mst010.

60. Steenwyk, J.L., Buida, T.J. III, Li, Y., Shen, X.-X., and Rokas, A. (2020). ClipKIT: A multiple sequence alignment trimming software for accurate phylogenomic inference. PLOS Biol. 18, e3001007. 10.1371/journal.pbio.3001007.

61. Wong, T.K.F., Ly-Trong, N., Ren, H., Demotte, P., Baños, H., Roger, A.J., Susko, E., Bielow, C., De Maio, N., Goldman, N., et al. (2026). IQ-TREE 3: phylogenomic inference software using complex evolutionary models. Mol. Biol. Evol. 43, msag117. 10.1093/molbev/msag117.

62. Kalyaanamoorthy, S., Minh, B.Q., Wong, T.K.F., von Haeseler, A., and Jermiin, L.S. (2017). ModelFinder: fast model selection for accurate phylogenetic estimates. Nat. Methods 14, 587–589. 10.1038/nmeth.4285.

63. Minh, B.Q., Nguyen, M.A.T., and von Haeseler, A. (2013). Ultrafast Approximation for Phylogenetic Bootstrap. Mol. Biol. Evol. 30, 1188–1195. 10.1093/molbev/mst024.

64. Guindon, S., Dufayard, J.-F., Lefort, V., Anisimova, M., Hordijk, W., and Gascuel, O. (2010). New Algorithms and Methods to Estimate Maximum-Likelihood Phylogenies: Assessing the Performance of PhyML 3.0. Syst. Biol. 59, 307–321. 10.1093/sysbio/syq010.

65. Letunic, I., and Bork, P. (2024). Interactive Tree of Life (iTOL) v6: recent updates to the phylogenetic tree display and annotation tool. Nucleic Acids Res. 52, W78– W82. 10.1093/nar/gkae268.

66. Kropinski, A.M., Mazzocco, A., Waddell, T.E., Lingohr, E., and Johnson, R.P. (2009). Enumeration of Bacteriophages by Double Agar Overlay Plaque Assay. In: Clokie, M.R., Kropinski, A.M. (eds) Bacteriophages. Methods in Molecular Biology™, vol 501. (Humana Press), pp. 69–76. 10.1007/978-1-60327-164-6_7.

67. Datsenko, K.A., and Wanner, B.L. (2000). One-step inactivation of chromosomal genes in Escherichia coli K-12 using PCR products. Proc. Natl. Acad. Sci. 97, 6640–6645. 10.1073/pnas.120163297.

68. Sullivan, A.E., Nabhani, A., Izrailevsky, D.S., Schinkel, K., Hoffman, C.R.K., Robbins, L.K., Nagy, T.A., Duncan, M.L., Ledvina, H.E., Erbse, A.H., et al. (2025). The Panoptes system uses decoy cyclic nucleotides to defend against phage. Nature 647, 988–996. 10.1038/s41586-025-09557-z.

69. Punjani, A., Rubinstein, J.L., Fleet, D.J., and Brubaker, M.A. (2017). cryoSPARC: algorithms for rapid unsupervised cryo-EM structure determination. Nat. Methods 14, 290–296. 10.1038/nmeth.4169.

70. Abramson, J., Adler, J., Dunger, J., Evans, R., Green, T., Pritzel, A., Ronneberger, O., Willmore, L., Ballard, A.J., Bambrick, J., et al. (2024). Accurate structure prediction of biomolecular interactions with AlphaFold 3. Nature 630, 493–500. 10.1038/s41586-024-07487-w.

71. Meng, E.C., Goddard, T.D., Pettersen, E.F., Couch, G.S., Pearson, Z.J., Morris, J.H., and Ferrin, T.E. (2023). UCSF ChimeraX: Tools for structure building and analysis. Protein Sci. 32, e4792. 10.1002/pro.4792.

72. Emsley, P., Lohkamp, B., Scott, W.G., and Cowtan, K. (2010). Features and development of *Coot*. Acta Crystallogr. D Biol. Crystallogr. 66, 486–501. 10.1107/S0907444910007493.

73. Liebschner, D., Afonine, P.V., Baker, M.L., Bunkóczi, G., Chen, V.B., Croll, T.I., Hintze, B., Hung, L.-W., Jain, S., McCoy, A.J., et al. (2019). Macromolecular structure determination using X-rays, neutrons and electrons: recent developments in *Phenix*. Acta Crystallogr. Sect. Struct. Biol. 75, 861–877. 10.1107/S2059798319011471.

74. Stephen, C., and Mishanina, T.V. (2022). Alkaline pH has an unexpected effect on transcriptional pausing during synthesis of the Escherichia coli pH-responsive riboswitch. J. Biol. Chem. 298, 102302. 10.1016/j.jbc.2022.102302.

75. Saikrishnan, K., Yeeles, J.T., Gilhooly, N.S., Krajewski, W.W., Dillingham, M.S., and Wigley, D.B. (2012). Insights into Chi recognition from the structure of an AddAB-type helicase–nuclease complex. EMBO J. 31, 1568–1578. 10.1038/emboj.2012.9.

76. Saikrishnan, K., Griffiths, S.P., Cook, N., Court, R., and Wigley, D.B. (2008). DNA binding to RecD: role of the 1B domain in SF1B helicase activity. EMBO J. 27, 2222–2229. 10.1038/emboj.2008.144.

77. Fang, C., Li, L., Shen, L., Shi, J., Wang, S., Feng, Y., and Zhang, Y. (2019). Structures and mechanism of transcription initiation by bacterial ECF factors. Nucleic Acids Res. 47, 7094–7104. 10.1093/nar/gkz470.

